# *In situ* electro-sequencing in three-dimensional tissues

**DOI:** 10.1101/2021.04.22.440941

**Authors:** Qiang Li, Zuwan Lin, Ren Liu, Xin Tang, Jiahao Huang, Yichun He, Haowen Zhou, Hao Sheng, Hailing Shi, Xiao Wang, Jia Liu

**Affiliations:** John A. Paulson School of Engineering and Applied Sciences, Harvard University, Cambridge, MA, USA; Broad Institute of MIT and Harvard, Cambridge, MA, USA; Department of Chemistry and Chemical Biology, Harvard University, Cambridge, MA, USA; Department of Chemistry, Massachusetts Institute of Technology, Cambridge, MA, USA

**Author notes:** These authors contributed equally (Qiang Li, Zuwan Lin, Ren Liu, Xin Tang).

## Abstract

Pairwise mapping of single-cell gene expression and electrophysiology in intact three-dimensional (3D) tissues is crucial for studying electrogenic organs (*e.g.*, brain and heart)^1–5^. Here, we introduce *in situ* electro-sequencing (electro-seq), combining soft bioelectronics with *in situ* RNA sequencing to stably map millisecond-timescale cellular electrophysiology and simultaneously profile a large number of genes at single-cell level across 3D tissues. We applied *in situ* electro-seq to 3D human induced pluripotent stem cell-derived cardiomyocyte (hiPSC-CM) patches, precisely registering the CM gene expression with electrophysiology at single-cell level, enabling multimodal *in situ* analysis. Such multimodal data integration substantially improved the dissection of cell types and the reconstruction of developmental trajectory from spatially heterogeneous tissues. Using machine learning (ML)-based cross-modal analysis, *in situ* electro-seq identified the gene-to-electrophysiology relationship over the time course of cardiac maturation. Further leveraging such a relationship to train a coupled autoencoder, we demonstrated the prediction of single-cell gene expression profile evolution using long-term electrical measurement from the same cardiac patch or 3D millimeter-scale cardiac organoids. As exemplified by cardiac tissue maturation, *in situ* electro-seq will be broadly applicable to create spatiotemporal multimodal maps and predictive models in electrogenic organs, allowing discovery of cell types and gene programs responsible for electrophysiological function and dysfunction.

## Main text

Simultaneously charting single-cell gene expression and electrophysiology in intact three-dimensional (3D) tissues across time and space is crucial to understanding the gene-to-function relationship in fields ranging from developmental biology to cardiology and neuroscience^1–5^. Such multimodal methods require stable and continuous recording of individual cell electrical activity with high spatiotemporal resolution across 3D tissue, multiplexed profiling of a large number of genes in electrically recorded cells, and cross-modal computational analysis.

Large-scale single-cell electrical recording^6–8^ and high-throughput single-cell sequencing^9–11^ have enabled system-level investigation of single-cell electrophysiology and gene expression, respectively. However, existing multimodal methods either lack high spatiotemporal resolution across the 3D tissue or cannot simultaneously measure tissue-wide electrical activities in a long-term stable manner. For example, combining calcium imaging with RNA hybridization^12^ can reveal the correlation of calcium activity and molecularly defined cell types. It, however, can only record the cell activity at second-timescale and profile a limited number of genes. Patch-seq^2, 3^ quantifies cell activity at millisecond temporal resolution and profiles the whole transcriptomes of the recorded cells but assays cells one at a time and requires membrane disruption during the electrical measurement, which is a challenge to tissue-wide and long-term stable electrical activity mapping.

Recent developments in thin-film flexible bioelectronics have enabled soft “tissue-like” electronics, capable of seamlessly integrating with tissue networks for long-term stable, millisecond-timescale single-cell electrical mapping^13–17^. Meanwhile, current imaging-based *in situ* sequencing methods^18^ can achieve end-point spatial analysis of thousands of genes at subcellular resolution across intact 3D tissues. Here, we integrate soft “tissue-like” bioelectronics with *in situ* sequencing as one method termed “*in situ* electro-seq” to enable a scalable and simultaneous profiling of single-cell electrophysiology and gene expression in intact 3D tissues.

### *In situ* electro-seq platform

We applied stretchable mesh electronics with cellular size electrodes to seamlessly integrate with tissue for continuous recording and to prevent potential cell-to-sensor dislocation from the tissue development, sample preparation, or multiple cycles of *in situ* sequencing (Fig. 1a). Then, we embedded the tissue-electronics hybrid into a hydrogel to form a tissue-electronics-hydrogel network, compatible with *in situ* sequencing protocol and capable of co-deformation during volume change. Meanwhile, to precisely identify the electrically recorded cells in 3D tissues, we used photolithography to pattern the thin-film microscale polymeric structures with unique fluorescent electronic barcodes (E-barcodes), paired with each electrode to label and locate their recording channel during fluorescence imaging cycles of *in situ* sequencing.

**Fig. 1.**
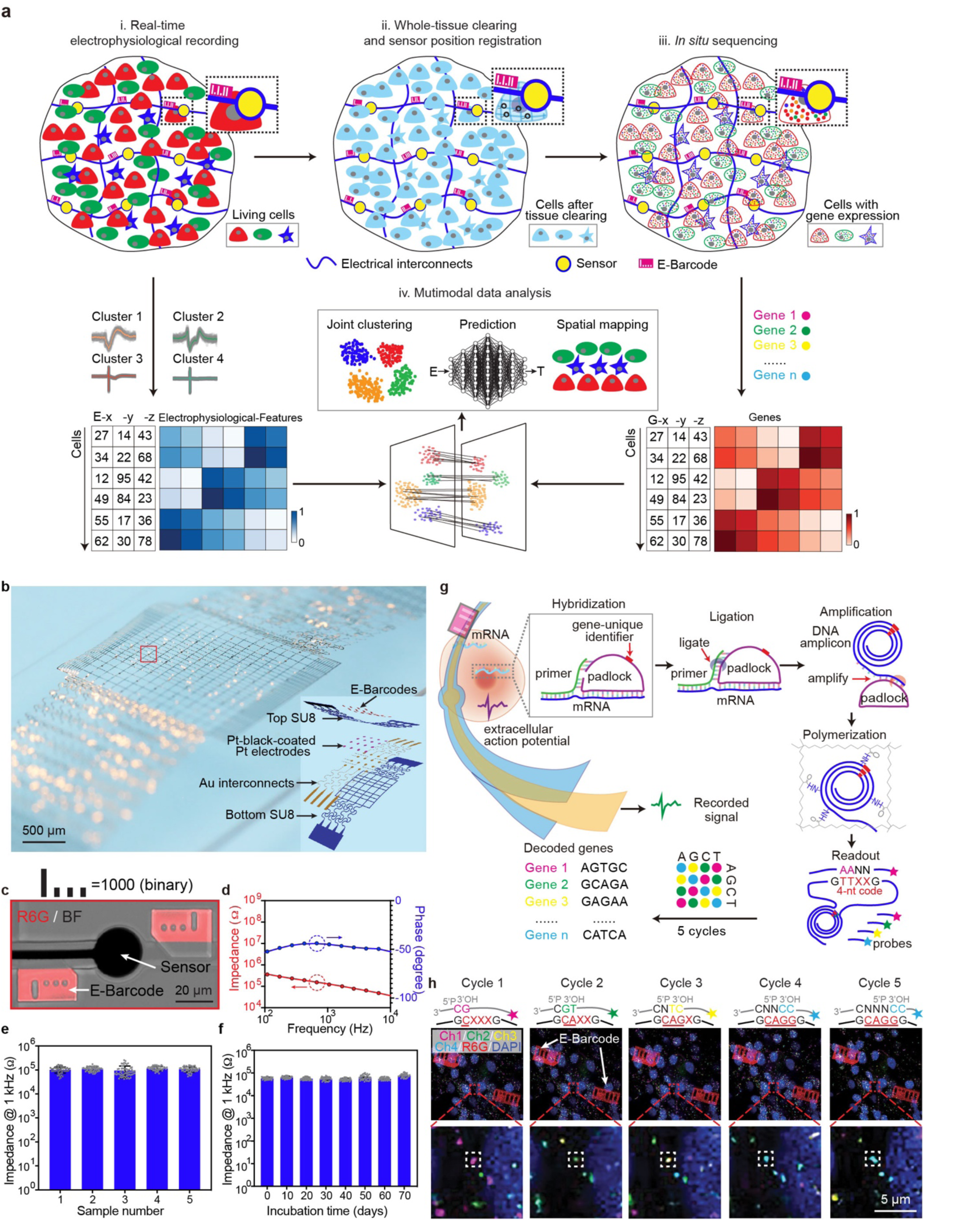
*In situ* electro-seq platform. **a**, Schematic summary of *in situ* electro-seq: integration of three-dimensional (3D) *in situ* sequencing and single-cell electrophysiology in tissue network. The flexible macroporous electronics are embedded across a 3D tissue network for continuous electrical recording at single-cell resolution. After electrical recording, the tissue-electronics hybrid is fixed for the *in situ* sequencing to read out the spatially-resolved gene expression information for each cell. Each electronic sensor is labeled by a fluorescent electronic barcode (E-barcode) defined by lithographic patterning (highlighted in dash boxes) to spatially register the sensor-to-cell position during imaging, which integrates the electrical recording and gene expression at the single-cell level. The integrated electrical and gene expression mapping is used for multimodal analysis such as joint clustering and cross-modal prediction. **b**, Representative photograph of the flexible mesh electronics for cardiac patch integration. Inset: schematic illustrates the multilayer structure of flexible mesh electronics. **c**, Overlap of fluorescence and bright-field (BF) images shows a pair of binary E-barcodes that label one electrode channel from the red box highlighted region in (**b**). **d**, Impedance and phase from 0.1 to 10 kHz of one representative electrode. (**e-f**), Averaged electrochemical impedance of electrodes across five representative mesh electronics at 1 kHz (**e**) and averaged impedance of electrodes at 1 kHz as a function of incubation time in PBS at 37°C from one representative stretchable electronics (n=64 electrode for each device, value=mean ± s.e.m., two-tailed, unpaired, t-test) (**f**). **g**, *In situ* sequencing of tissue-electronics hybrid. The custom padlock probe and primer hybridize to intracellular mRNA of the 3D tissue-electronics hybrid, followed by enzymatic ligation and rolling circle amplification (RCA) to construct the *in situ* cDNA amplicons. The amplicons are then copolymerized with acrylamide, forming the hydrogel network. A gene-specific identifier (red) in the probe is amplified through this process, which could then be used for sequencing. After *in situ* sequencing of cDNA amplicons in tissue-electronics-hydrogel network, the gene-specific identifier is decoded through multiple sequencing cycles. **h**, Representative images of five rounds sequencing and co-imaging of E-barcode. In each sequencing cycle, the reading probe has one increasing number of degenerative base N which sets off the starting position for sequencing at 5’phosphate; the decoding probes (gray line on the right with star-symbol label) are color coded with fluorophore at the 3’ end according to the 2-base encoding diagram. Both reading probe and decoding probe hybridize to the gene-specific identifier followed by ligation and imaging. After imaging, both reading and decoding probes are stripped away with 60% formamide. X, unknown base; red underline, decoded sequence; Ch1 to Ch4, fluorescence channels; E-barcodes labelled with R6G.

*In situ* electro-seq consists of the following four key steps (Fig. 1a): (i) the mesh electronics with E-barcoded sensors are embedded in tissues for continuous single-cell electrical recording; (ii) the entire tissue-electronics hybrid is fixed, embedded in hydrogels, and cleared for *in situ* sequencing; (iii) gene identities and E-barcodes are simultaneously read out by multiple cycles of fluorescence imaging, integrating electrical recording with gene expression profiling at single-cell resolution; and (iv) the integrative data are analyzed using multimodal and cross-modal visualization, correlation and prediction to illustrate the spatiotemporal gene-to-electrophysiology relationship. As a demonstration, we applied *in situ* electro-seq to a human induced pluripotent stem cell-derived cardiomyocyte (hiPSC-CM) patch, simultaneously mapping its electrophysiology and gene expression. Figure 1b shows a representative mesh electronics with 64 electrodes with 25-µm diameter (Fig. 1b and Extended Data Fig. 1a-e), which is similar to a typical size of the CM^19^, and enables single-cell electrophysiological recordings. A pair of center-symmetric fluorescent E-barcodes with unique binary codes were patterned with each electrode as center (Fig. 1c and Extended Data Fig 1f-g). Characterization of electrode impedances (Fig. 1d) showed stable electrochemical performance across different samples (Fig. 1e) and over >2 months incubation in the physiological solution (Fig. 1f) for long-term electrical recording.

HiPSC-CMs were cultured with mesh electronics on a Matrigel layer to form a 3D cardiac patch as described in previously reported methods^15, 17^ (Extended Data Fig. 2a-e, see Methods). After culturing cells with mesh electronics and recording, we fixed the tissue-electronics hybrid and applied *in situ* sequencing protocol (see Methods) to profile a targeted set of cardiac genes, including the 201 most differentially expressed genes during cardiac maturation extracted from published single-cell RNA sequencing (scRNA-seq) data^20, 21^. RNA-derived DNA amplicons with pre-designed gene-specific identifiers were synthesized *in situ* by probe hybridization, enzymatic amplification, and immobilization in the cleared tissue-electronics-hydrogel network (Fig. 1g). Then, the gene-specific identifier was decoded through five sequencing cycles^18^. Notably, the microscale distances among E-barcodes, cells, and amplicons remained during the multiple cycle imaging (Fig. 1h).

### *In situ* electro-seq enabled correlation of electrophysiological and transcriptional data

We first tested *in situ* electro-seq to the hiPSC-CM cardiac patch at Day 46 of differentiation (Fig. 2a-d). 64-channel recording was applied with their single-spike action potential waveforms identified, pre-processed, averaged and down-sampled to extract 62 features (see Methods). Figure 2e and f show the representative 16-channel voltage traces and single-spiked waveforms, respectively. The uniform manifold approximation and projection (UMAP)^22^ visualization (Fig. 2f inset) of extracted features from 64 channels shows the heterogeneity of hiPSC-CM electrophysiology. *In situ* sequencing was applied immediately after electrical recording (Fig. 2g, Extended Data Fig. 3a). After 3D cell segmentation (ClusterMap^23^, Fig. 2h-i), we performed cell clustering using 201 genes from 32,429 cells across all the imaged positions. Leiden clustering^24^ showed two major cell types (Fig. 2j), CMs and fibroblasts (Fibs), which were spatially mapped back to E-barcoded electrodes (Fig. 2k). We built a computational pipeline to automatically identify CMs that formed directly contacted with electrodes as electrically recorded cells (Fig. 2k and Extended Data Fig. 3b). Then, identification of E-barcodes registered the electrophysiological features with gene expression of the electrically recorded cells. Heatmap (Fig. 2l) and joint UMAP (Fig. 2m) visualization showed the integrative z-scored electrophysiological features and 24 top-ranked differentially expressed CM-related genes, and their multimodal distributions at single-cell level, respectively.

**Fig. 2.**
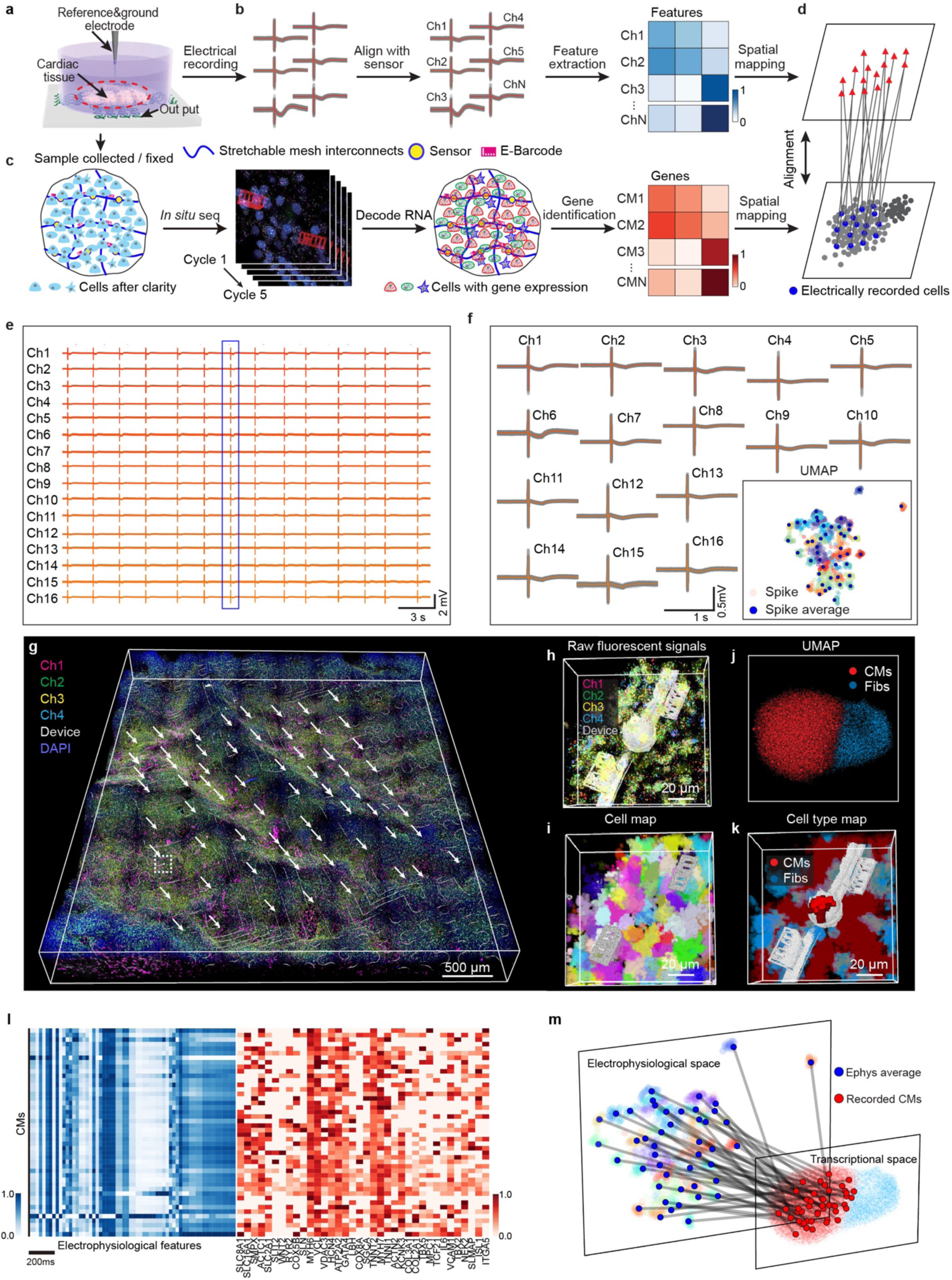
*In situ* electro-seq simultaneously maps single-cell transcriptional and electrophysiological states of human cardiac patch. **a-d**, Schematics of experiment: Flexible mesh electronics with 64-channel subcellular size electrode array is embedded with human induced pluripotent stem cell-derived cardiomyocyte (hiPSC-CM) patch for continuous electrical recording (**a**). *In situ* electro-seq records the extracellular action potential (**b**), profiles spatially-resolved RNA expression (**c**), and integrates multimodal data at the single-cell level (**d**). **e**, Representative 16-channel voltage traces recorded from cardiac patch at Day 46 of differentiation. **f**, Representative 16-channel averaged single spike waveform of action potential detected from 1-min recording from (**e**). Inset shows the uniform manifold approximation and projection (UMAP) visualization of the spike waveforms from 64 channels. Blue dots highlight the distribution of the averaged spike waveform for each channel. **g**, Raw fluorescence imaging of in-process *in situ* electro-seq for 201 genes with the full view of the entire tissue-electronics hybrid. White arrows highlight positions of 64 electrodes. **h**, Zoomed-in view of the raw fluorescent signals illustrates the representative electrode embedded area from the white dashed box highlighted regions in (**g**). **i**, 3D cell segmentation map generated by ClusterMap labels cells with different colors. **j**, The UMAP visualization represents major cell types across 32,429 cells in the entire cardiac patch clustered by Leiden clustering. **k**, 3D cell-type map labels each cell by its cell type with the same color code in (**j**). The electrically recorded cell is highlighted by deep red. **l**, Heatmap shows the extracted features from the waveform of averaged spikes and corresponding 36 top differentially expressed genes expressed in the electrically recorded cells (see Methods). Colors correspond to normalized electrophysiological feature value (blue) and gene expression value (red). **m**, Integration of electrophysiological with gene expression features for each cell in UMAP visualization by identifying electrode-to-cell positions through imaging of E-barcodes.

### *In situ* electro-seq of cardiac maturation

We used *in situ* electro-seq to trace the maturation of the hiPSC-CM patch (Fig. 3a). The cell electrophysiological signals from Day 12, Day 21, Day 46, and Day 64 of differentiation (Extended Data Fig 4) showed the distinct features at these four stages (Fig. 3a). We applied our recently developed ClusterMap^23^ method to segment cells by RNA amplicons, and compared with the conventional cell nuclei-based segmentation method StarDist^25^, ClusterMap can identify higher RNA counts-per-cell across samples robustly with less batch variation across different differentiation days (Extended Data Fig. 5) and showed the similar cell-typing results as StarDist (Extended Data Fig. 6). Comparing UMAP visualization of cell types across four differentiation stages showed similar embedding distributions between electronics-embedded and control samples, which suggests negligible effects of mesh electronics on cardiac maturation and performance of *in situ* sequencing (Extended Data Fig 6a-b). We performed unsupervised Leiden clustering analysis on all of the *in situ* sequenced cells (130,162 cells) across 4 stages, which identified 5 cell clusters (Extended Data Fig 6c). Based on the expression profiles of the marker genes, four subpopulations can be attributed to CMs and one to the Fibs. Compared with previous reports^26, 27^, the expression level of marker genes (*e.g.*, HCN4, MYH6, MYH7, MYL7, MYL4, *etc*.) in these four types of CMs indicates the transition of cardiac states from nodal-like through atrial-like CMs to ventricular-like CMs (Extended Data Fig 6d-e), which can be further confirmed by the gene expression profiles over time--the nodal marker gene (HCN4) and atrial marker gene (MYH6) were decreased at the later stage while the ventricular maker gene (MYH7) is increased (Extended Data Fig 6f-g). Gene ontology (GO) analysis of the top differentially expressed genes revealed enriched gene sets in cardiac muscle contraction, ion conduction, and tissue development (Extended Data Fig 6h). Collectively, these data demonstrate that the *in situ* electro-seq protocol can identify the evolution of electrophysiological and transcriptional profiles over the time course of hiPSC-CM maturation.

**Fig. 3.**
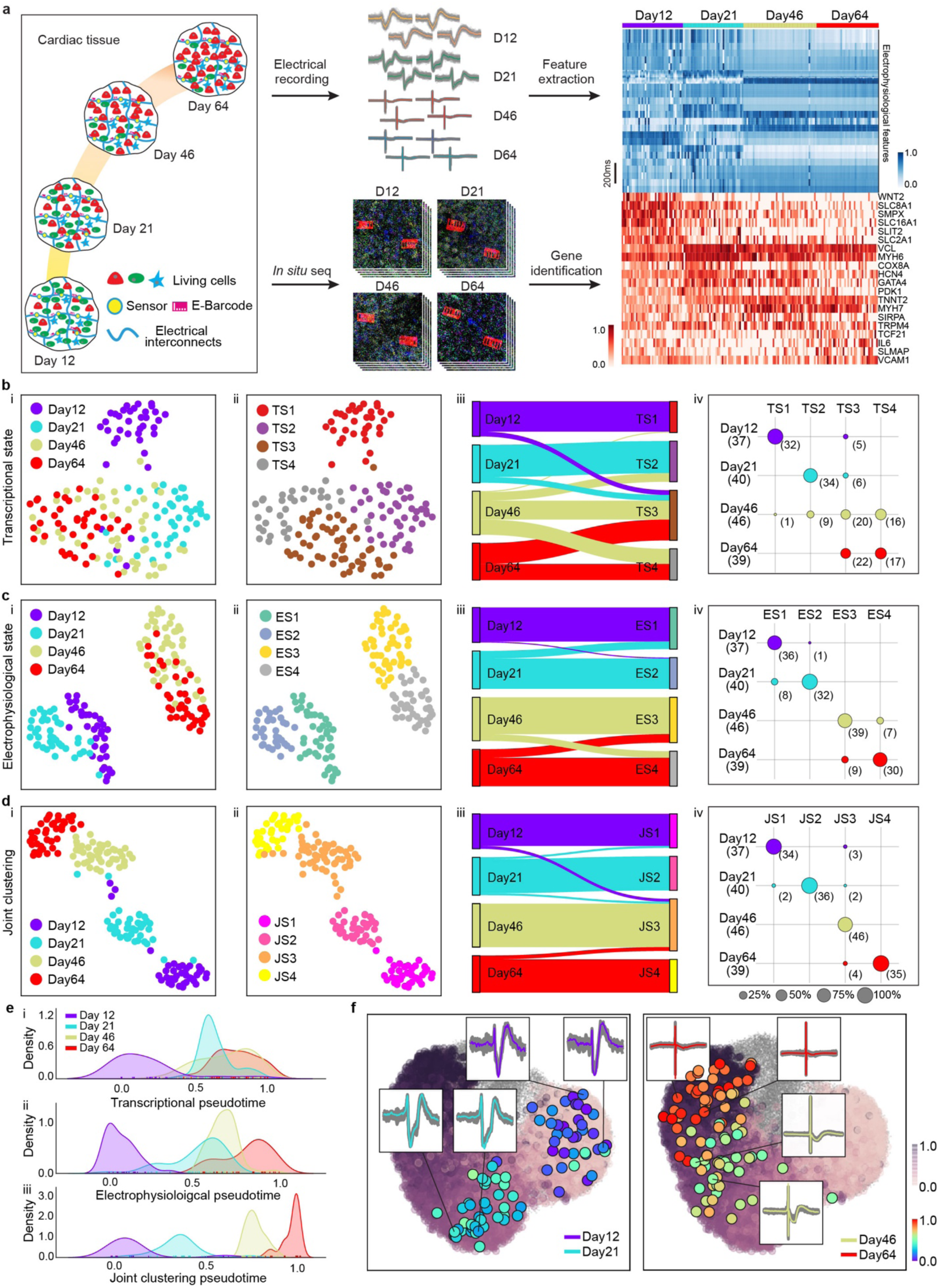
*In situ* electro-seq enables joint clustering of cell states in 3D human cardiac tissue maturation. **a,** Overview schematics illustrate the application of *in situ* electro-seq to 3D hiPSC-CM patches at different stages. hiPSC-CM patches are integrated with flexible mesh electronics and fixed for *in situ* electro-seq at Day 12, Day 21, Day 46, and Day 64 of differentiation. Integrated heatmap plots electrophysiological features from 162 cells across 4 samples and their corresponding 20 top differentially expressed gene expression profiles. **b**, hiPSC-CM transcriptional states (t-states, TS) defined by gene expression. UMAP visualization of gene expression of electrically recorded CMs that are color coded by differentiation days (**i**) and t-states defined by Leiden clustering (**ii**). Comparison of t-states and differentiation days by river plot (**iii**) and dot plot (**iv**). Size of dots represents the percentages of cells from given differentiation days that match to the corresponding t-states. Each row sums to 100%. **c-d**, hiPSC-CM electrophysiological states (e-states, ES) (**c**) and transcriptional/electrophysiological joint states (j-states, JS) defined by Weighted Nearest Neighbor (WNN)-integrated representations from gene expression and electrophysiology (**d**) are analyzed as (**b**). **e**, Distribution plots show pseudotime distributions of all the electrically recorded CMs that are learned from *Monocle3* using gene expression (**i**), electrophysiology (**ii**), and WNN-integrated representations of gene expression and electrophysiology (**iii**). The pseudotime analyses are normalized to 0-1. **f**, Electrically recorded cells with colors encoding the joint pseudotime in (**e**) are mapped to the UMAP visualization of gene expression from all 130,162 cells sequenced from samples across four stages that are shown as gray embedding. All 112,892 CMs from these four stages are labeled with purple colors encoding gene expression defined pseudotime. Insets show representative single-spike waveforms from Day 12, Day 21, Day 46, and Day 64 of differentiation.

### *In situ* electro-seq-enabled multimodal joint analysis

*In situ* electro-seq is capable of integrating gene expression with electrophysiology for joint cell clustering. While gene expression clustering of *in situ* sequenced 112,892 CMs can separate CMs into four transcriptional states (t-states), roughly corresponding to samples collected at the four differentiation days (Extended Data Fig. 6c), we noticed that the t-states of CMs at Day 46 and 64 of differentiation are less separable (Extended Data Fig. 7d, i). The silhouette score (s-score) matrix shows that the separability of t-states of 162 electrically recorded CMs is worse (Fig. 3b, Extended Data Fig. 7a-c), especially at Day 21, Day 46 and Day 64 of differentiation (Extended Data Fig. 7d, ii), which agrees with previously reported scRNA-seq clustering of hiPSC-CMs t-states during maturation^27^. Using electrophysiological features to cluster the electrically recorded cells, two major electrophysiological states (e-states) were identified for CMs, one for Day 12 and 21, and the other for Day 46 and 64, respectively. However, the separability within the two major groups is low (Fig. 3c, Extended Data Fig. 7d, iii). Previous patch-seq results suggest that integration of gene expression with electrophysiology can improve the classification of cell types^28^. Here, we used the weight nearest neighbor (WNN)^29^ algorithm from *Seurat* v4 to integrate the single-cell electrophysiological and gene expression data from *in situ* electro-seq for joint representation (see Methods). Using the joint features, CMs can be clustered into four joint states (j-states) that are well separated (Fig. 3d, Extended Data Fig. 7d, iv) and better represent the different differentiation days than both t-states and e-states quantified by the improved matching accuracy (Fig. 3b-d, iii-iv).

Furthermore, to test whether the multimodal data integration and joint representation can also better resolve continuous CM trajectory during maturation, we applied *Monocle3*^30^, an unsupervised method, to calculate the pseudotime distributions of t-states, e-states, and j-states. The results also show that the integrated gene expression and electrophysiology data improved the separation of pseudotime distributions for cells at different differentiation stages (Fig. 3e). Highlighting the 162 electrically recorded CMs with their j-state pseudotimes in the UMAP visualization of total 112,892 CMs t-states (Fig. 3f) shows that, with as few as 162 cells, the j-states clearly recapitulated the continuous maturation trajectory of 112,892 CMs (Extended Data Fig. 6f).

### *In situ* electro-seq-enabled cross-modal correlation, prediction and mapping

A unique capability of *in situ* electro-seq is to use continuous single-cell electrical measurement to predict and infer a continuous mapping of gene expression from the same tissue during development and function. To examine to what extent the t-state can be predicted by e-state during cardiac maturation, we applied sparse reduced-rank regression (RRR)^31^ to visualize the gene-to-electrophysiology relationship. The 62 electrophysiological features from each cell were used (Fig. 4a and Extended Data Fig. 8e). Cross-validation was performed by tuning the regularization strength (Extended Data Fig. 8a-d). The selected model chose 32 genes with a 5-dimensional latent space and achieved a cross-validated *R*^2^ of 0.2 for CM-correlated genes. Then, to visualize the structure of the latent space, we projected gene expression and electrophysiological features onto the latent dimensions (Extended Data Fig. 8f). The cross-validated correlation between the first two pairs of projections was 0.72 and 0.49, respectively. These first two components separate CM groups by their days of differentiation. These different groups show distinct correlated genes with electrophysiology features. The model detected cardiac structural related genes such as myosin heavy chain (MYH6, MYH7), troponin complex (TNNT2), and Z-disc (VCL, VIM) as well as calcium signaling gene (RYR2) and mitochondrial genes (COX5B, COX8A) (Fig. 4c and Extended Data Fig. 8f). In addition, an RRR model restricted to using only ion channel genes also achieved *R*^2^=0.15, and correlations of 0.68 and 0.37 in the first two pairs of projection (Fig. 4b). Genes detected by the ion-channel-only model are endoplasmic reticulum calcium transporting gene (ATP2A2), sodium-calcium exchanger (SLC8A1), potassium channel (KCNK6, KCNQ1, KCND3), cyclic nucleotide-gated ion channel (HCN4) (Fig. 4b-c), whose molecular functions are consistent with the previous knowledge of action potential waveform^32, 33^.

**Fig. 4.**
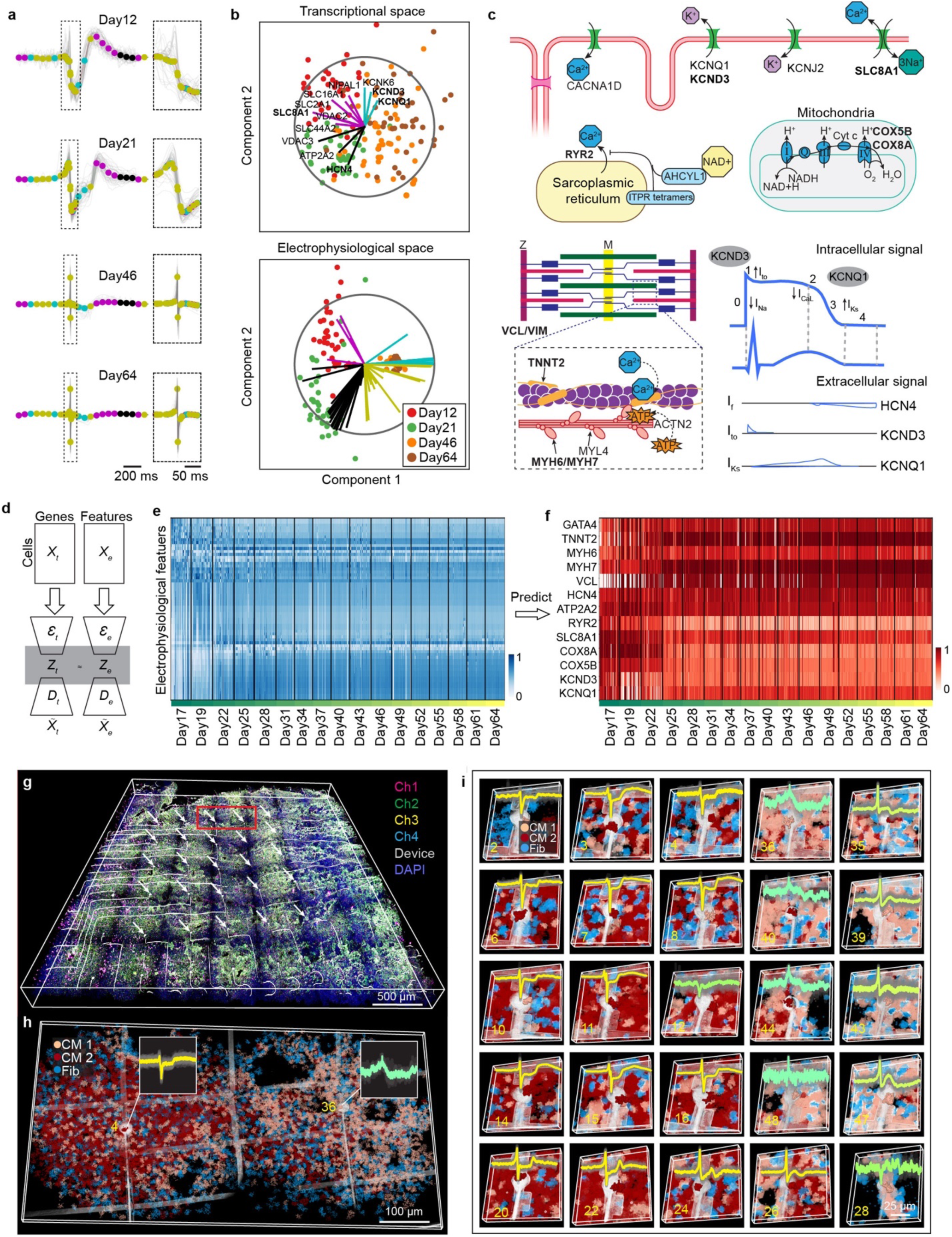
*In situ* electro-seq enables cross-modal visualization, correlation, prediction, and mapping. **a**, Representative 62 electrophysiological features are extracted through down sampling of each spike waveform on Day 12, Day 21, Day 46, and Day 64 of differentiation, respectively. 1.6-second waveforms are sampled to 20 bins. Inset: 0.15-second fast spikes are sampled to 42 bins. **b,** A sparse reduced-rank regression (RRR) model to visualize and align t-states and e-states. Components 1 and 2 of the rank-5 model are shown, n=162. The model selects 40 and 12 ion channel-related genes to train the model for visualization. **c**, Schematics show physiological functions of genes that are mostly correlated to electrophysiological feature changes during differentiation identified by the RRR model in (**a-b**) and **Extended Data Fig. 8e-f**. **d**, Schematics show the structure of a coupled autoencoder for electrophysiology-to-transcripts (E-to-T) prediction. Encoders (*Ɛ*) compress input data (*X*) from E and T modality into low-dimensional representations (*Z*), whereas decoders (*D*) reconstruct data (*X̅*) from representations **e**, Heatmap shows single-cell electrophysiological features continuously recorded from the same cardiac patch over the time course of maturation. **f**, Heatmap shows the single-cell gene expression profiles predicted by coupled autoencoder (**d**) using data in (**e**). The 13 electrophysiology-related genes shown here are selected from the differentiation-related gene sets identified by RRR models. The values are normalized to 0 to 1 for each electrical feature (**e**) and gene (**f**), respectively. **g**, *In situ* electro-seq enables the multimodal spatial mapping of the cardiac tissue with heterogeneous cell populations. **h**, Zoom-in image of the red box highlighted region in (**g**) shows the representative region that contains multiple, spatially-arranged cell populations with distinct electrophysiological activities (inset). **i**, 25 representative regions mapped by *in situ* electro-seq highlight spatially-solved cell types and electrophysiological waveforms from the white arrows labeled regions in (**g**). Color codes label different t-states and e-states for cells. Numbers indicate E-barcoded electrode channels.

To enable electrophysiology-to-gene prediction, we constructed a coupled autoencoder^34, 35^ to learn coordinated representations of integrative electrophysiology and gene expression data from Day 12, Day 21, Day 46, and Day 64 of differentiation generated by *in situ* electro-seq (Fig. 4d). Aligned 2D representations *Zt* and *Ze* for the high-dimensional gene expression and electrophysiology data *Xt* and *Xe* encoded from the encoder *Ɛ* (Extended Data Fig. 9a-b) showed the well separated and aligned gene expression and electrophysiology distributions. This result suggests a common latent representation exists across gene expression and electrophysiological data so that *in situ* electro-seq-generated electrophysiology data can be used to predict gene expression^34, 35^. We then applied the coupled autoencoder to the electrical measurement recorded every three days of a hiPSC-CM patch over the time course of maturation from Day 17 to Day 64 of differentiation (Fig. 4e and Extended Data Fig. 9c-d). The predicted single-cell gene expression profile shows the evolution of electrophysiology-related genes identified by RRR models (Fig. 4f, Extended Data Fig. 9e-f). We further applied the coupled autoencoder to the electrical measurement of a millimeter-scale hiPSC-CM organoid with mesh electronics fully embedded across the 3D volume (Extended Data Fig. 9g-j). The predicted single-cell gene expression profile shows the evolution of electrophysiology-related genes with much higher variation at the single-cell level (Extended Data Fig. 9k-l), suggesting the intrinsic heterogeneity in hiPSC-CM maturation in 3D organoids^33^.

We applied the *in situ* electro-seq to the heterogeneous cardiac patch formed by hiPSC-CMs at different maturation stages (Fig.4g and Extended Data Fig. 10). The results showed accurate mapping of the gene expression and electrophysiology heterogeneity at single-cell resolution from the sample (Fig. 4h-i), suggesting its applicability to the tissues with spatially heterogeneous cell types and states.

## Discussion

We have demonstrated that *in situ* electro-seq is capable of integrating electrophysiology and gene expression at the single-cell level, providing (i) multimodal joint cell clustering in spatially heterogenous tissues at single-cell resolution across different stages of hiPSC-CM maturation, which are challenging to be directly traced by previous approaches; (ii) cross-modal correlation and prediction that use continuous electrical measurement to predict and infer single-cell gene expression evolution from tissues; and (iii) identification of gene programs directly relevant to functional maturation. Future work may address potential limitations and opportunities; for example, different flexible nanoelectronic sensors and stimulators could integrate long-term stable intracellular electrophysiology^36^ and multifunctional (*e.g.*, electrical, mechanical, optical, chemical, *etc*.) single-cell recording and stimulation^37^ with *in situ* sequencing of different biomolecules (*e.g.*, RNA or DNA). Further integration of complementary metal-oxide-semiconductor multiplexing circuits could also substantially increase the number of cells that can be simultaneously measured^6^. The *in situ* electro-seq may be applied to *in vivo* tissues from behaving animals to map single-cell gene expression and functions from cardiac and neural systems as well as their pathophysiological states in which tissue-wide electrophysiological dysfunctions are related to the cell-level gene expression variations such as neuropsychiatric diseases^38, 39^ (*e.g.*, autism spectrum disorder, bipolar disorders, *etc.*) and cardiac arrhythmia^40^ (*e.g.*, atrial fibrillation, ventricular tachycardia, *etc.*).

## Methods

### 1. Stretchable mesh nanoelectronics

#### Fabrication of stretchable mesh electrode array

Fabrication of the ultra-flexible, stretchable mesh nanoelectronics was based on methods described previously^14–16^. Key steps are described as follows: 4-inch glass wafers (Soda lime glass) were used as a transparent and insulating substrate for fabrication and cell culture. The glass wafers were cleaned by piranha solution (3:1 mixture of sulfuric acid and 30% hydrogen peroxide), followed by rinsing with deionized (DI) water and drying with the N_2_. Hexamethyldisilazane (HMDS, MicroChem) was spin-coated at 4000 rpm to increase adhesion of photoresists with the substrate. LOR 3A (300 nm, MicroChem)/S1805 (500 nm, MicroChem) were spin-coated at 4000 rpm/4000 rpm, followed by baking at 180 °C for 5 mins and at 115 °C for 1 min, respectively. Ni sacrificial layer was exposed by using a Karl Suss MA6 mask aligner with 365 nm ultraviolet (UV) light at 40 mJ/cm^2^ and developed by CD-26 developer (MICROPOSIT) for 70 s. O_2_ plasma (Anatech Barrel Plasma System) was used for the removal of photoresist residues at 50 W for 30 s. Sharon Thermal Evaporator was used for the deposition of 100-nm-thick Ni followed by a standard lift-off procedure in remover PG (MicroChem) for 2 hours. After patterning the Ni layer, SU-8 precursor (SU-8 2000.5, MicroChem) was spin-coated at 4000 rpm, pre-baked at 65 °C / 95 °C for 2 mins each, exposed to 365 nm UV at 200 mJ/cm^2^, post-baked at 65 °C / 95 °C for 2 mins each, developed using SU-8 developer (MicroChem) for 60 s, rinsed by isopropyl alcohol (IPA) for 30 s, blow for drying by N_2_, and hard-baked at 180 °C for 40 mins to define mesh-like SU-8 400-nm-thick patterns as the bottom encapsulation layer. After patterning the SU-8 bottom layer, HMDS/LOR3A/S1805 photoresist layers were spin-coated as described above, followed by depositing 5/40/5-nm-thick chromium/gold/chromium (Cr/Au/Cr) by the electron-beam evaporator (Denton), and the standard lift-off procedure in the remover PG (MicroChem) overnight to define the Cr/Au interconnects. Then, the same photolithography process was used to define 5/50-nm-thick chromium/platinum (Cr/Pt) as electrodes. After patterning electrodes, the top SU-8 encapsulating layer was patterned using the same method described for patterning the bottom SU-8 layer. Finally, fluorescent E-barcodes were defined by patterning the SU-8 structure doped by adding 0.004 wt‰ of Rhodamine 6G powder (Sigma-Aldrich) into SU-8 precursor.

#### Connection of stretchable mesh electrode array with flexible cable for electrical recording

Next, the flexible flat cable (FFC, Molex) was soldered onto the input/output pads using a flip-chip bonder (Finetech Fineplacer), followed by gluing a culture chamber onto the substrate wafer to completely enclose the mesh part of the device using a biocompatible adhesive (Kwik-Sil, WPI). Then, Pt black (PtB) was electroplated on the Pt electrode array using a precursor of 0.08 wt% chloroplatinic acid (H_2_PtCl_6_) solution (Sigma-Aldrich) in H_2_O. The precursor was drop-casted onto the device, followed by passage of a 1 mA/cm^2^ DC electric current density for 3 mins using mesh electrodes as anodes and an external Pt wire as the cathode. The device was then rinsed with DI water for 30 s and dried by N_2_. Finally, the surface of the device was treated with oxygen plasma (Anatech 106 oxygen plasma barrel asher), followed by adding 1 mL of Ni etchant (type TFG, Transene) into the chamber for 2 to 4 hours to completely release the mesh electronics from the glass substrate. The device was then ready for subsequent sterilization steps before cell culture.

#### Electrochemical measurements

The electrochemical impedance spectra (EIS) of the electrodes were measured based on methods described previously^41^. The three-electrodes setup was used to measure the EIS of each electrode. A standard silver/silver chloride (Ag/AgCl) electrode and platinum wire (300 µm in diameter, 1.5 cm in length immersed) were used as reference electrode and counter electrode, respectively. The device was immersed in 1 X PBS solution (Thermofisher) during measurement. The SP-150 potentiostat (Bio-logic) along with its commercial software EC-lab was used to perform the measurements. For each measurement, at least three frequency sweeps were measured from 1 MHz down to 1 Hz to obtain statistical results. A sinusoidal voltage of 100 mV peak-to-peak was applied. For each data point, the response to 10 consecutive sinusoids (spaced out by 10% of the period duration) was accumulated and averaged.

### 2. Cell culture, tissue integration, and electrical recording

#### Cell culture and cardiomyocytes (CMs) differentiation

Human induced pluripotent stem cells (hiPSC, hiPSC-IMR90-1) were obtained from WiCell Research Institute (Madison, WI, USA). Authentication and testing for the mycoplasma were performed by WiCell Research Institute. hiPSC-IMR90-1 cells were cultured on a Matrigel-coated 6-well plate with Essential 8 medium (Gibco). The medium was changed daily. The cells were passaged every 3-4 days. hiPSC-derived cardiomyocytes were generated according to the methods described previously^26, 42^. The hiPSC-IMR90 cells were cultured on a Matrigel-coated 6-well plate with Essential 8 medium to 70-80% confluency before initiating cardiac differentiation. The first day was defined as day 0. For cardiac differentiation, the cells were maintained in RPMI 1640 medium (Gibco) plus 1% B27-insulin (Gibco). CHIR99021 (12 µM; BioVision) was applied on day 0; IWR1 (5 µM; Cayman) was applied from day 3 to day 4. The cardiac cells were maintained in RPMI 1640 medium plus 1% B27 (Gibco) from day 7 and the medium was changed every other day accordingly.

#### Integration of mesh electronics with hiPSC-CM patch

First, the released stretchable mesh electronics in the culture chamber was rinsed with DI water, decontaminated by 70% ethanol and incubated with Poly-D-lysine hydrobromide (0.1 mg/mL) overnight followed by coating with Matrigel solution (100 µg/mL) for about 1 hour at 37°C. Then, the device was pre-chilled on an ice bag in the biosafety hood and then 70 µL Matrigel solution (10 mg/mL) was added from the edge of the chamber to the cell culture medium, ensuring that the Matrigel covered the entire bottom substrate of the cell culture chamber underneath the stretchable mesh electronics. Next, the device was transferred into the incubator for at least 30 mins at 37 °C to cure the Matrigel solution into a Matrigel hydrogel layer. Finally, hiPSC-CMs were incubated with 0.05% Trypsin-EDTA solution (Biosciences) for 5 mins and then dissociated into single cells. About 3∼4 million cells were suspended in 1 mL RPMI 1640 medium plus 1% B27 and then transferred onto the cured electronics / Matrigel hybrids in the cell culture chamber and maintained at 37 °C, 5% CO_2_. 5 µM rock inhibitor (Y27632) was added to the medium in the first day to improve the cell viability. The CMs will form a continuous cell patch with the stretchable mesh electronics embedded within 24-48 hours. Notably, to include cells with different differentiation stages, ca. 2 million Day 18 of differentiation CMs were seeded from one side of the culture chamber and cultured for 5 days; then, another ca. 2 million Day 7 of differentiation CMs were seeded from the opposite side of the cell culture chamber.

#### Electrophysiological measurement

The Blackrock CerePlex Direct voltage amplifier along with a 32 channel Blackrock µ digital headstage connected to the device were used to record electrical activity from the cardiac patch. The culture medium was grounded by a Pt electrode. A second Pt electrode was used as a reference electrode. During electrical measurement, samples were placed on a battery powered warming plate that maintained thermostatic 37 ℃. The measurement setup was placed into a Faraday cage. A sampling rate of 30,000 Hz was used for the electrical recording. The cell electrical activities were recorded every 3 days. MATLAB and Python codes provided by Blackrock were used to load, view, and convert raw data files into an accessible format for data analysis.

### 3. *In situ* sequencing

*In situ* sequencing experiments were performed based on methods described previously with some modifications^18^. Glass-bottom 12-well plates (Mattek, P12G-1.5-14-F) were first treated with oxygen plasma for 5 mins (Anatech Barrel Plasma System, 100W, 40% O_2_) followed by methacryloxypropyltrimethoxysilane (Bind-Silane) solution (88% ethanol, 10% acetic acid, 1% Bind-Silane, 1% H_2_O) treatment for 1 hour. The 12-well plates were then rinsed with ethanol for 3 times and were left to dry at room temperature (R.T.) for 3 hours. The 12-well plates were further treated with 0.1 mg/mL of Poly-D-lysine solution for 1 hour at R.T. followed by 3 times rinsing with H_2_O. The plates were air-dried for 1 hour at R.T. Micro cover glasses (12 mm) were pretreated with Gel Slick at R.T. for 10 mins and were then air-dried before using.

The cardiac tissue was fixed with 1 mL 1.6% PFA for 30 mins at R.T. and then washed with PBS 3 times for 10 mins each time. The sample was then transferred from the chamber to the 12-well plates and permeabilized with 1mL (0.1M glycine, 0.1 U/µL SUPERase·In, 0.5% Triton-X 100 in PBS) for 30 mins. The sample was washed with 1mL PBST (0.1% Triton-X 100 in PBS) 3 times for 10 mins each. The sample was then incubated in 1X hybridization buffer (2X SSC, 10% formamide, 1% Triton-X 100, 20 mM RVC, 0.1 mg/mL yeast tRNA and pooled SNAIL probes at 20 nM per oligo) in a 40 °C humidified oven with gentle shaking for 48 hours. The sample was washed with 1ml PBSTV (1% RVC in PBST) at 37 °C 3 times for 20 mins each and washed with high salt buffer (4X SSC in PBST) for another 20 mins at 37 °C, and then washed with PBST three times for 10 mins each at R.T. The sample was then incubated in 1mL ligation mixture (1: 50 T4 DNA ligase, 1:100 BSA, 0.2 U/µL SUPERase-In) at R.T. overnight and then washed with 1mL PBST three times for 10 mins each. The sample was incubated in 1mL RCA mixture (1: 50 Phi29 DNA polymerase, 250 µM dNTP, 1:100 BSA, 0.2 U/µL SUPERase-In and 20 µM 5-(3-aminoallyl)-dUTP) at 4 °C for 1 hour before incubating at 30 °C for 6 hours and then washed with 1 mL PBST 3 times for 10 mins each. The sample was incubated with 20 mM acrylic acid NHS ester in PBST for 3 hours at R.T. and washed with PBST 3 times for 10 mins each. The sample was then incubated with monomer buffer (4% acrylamide, 0.2% bis-acrylamide, 2X SSC) overnight at R.T. The buffer was then aspirated and 55 µL of polymerization mixture (0.2% ammonium persulfate, 0.2% tetramethylethylenediamine dissolved in monomer buffer) was added to the sample. The Gel Slick coated coverslip was immediately put on the sample and the polymerization was conducted in an N_2_ container for 90 mins. The sample was then washed with PBST 3 times for 10 mins each.

Five cycles of sequencing experiments were performed to decode gene identity. Within each cycle, the sample was first treated with a stripping buffer (60% formamide, 0.1% Triton-X-100) at R.T. for 6 times, 15 mins each followed by PBST wash for 6 times, 10 mins each. Then the sample was incubated with the sequencing mixture (1: 25 T4 DNA ligase, 1: 100 BSA, 10 µM reading probe and 5 µM fluorescent oligos) at R.T. for 12 hours. Then the sample was washed by the washing and imaging buffer (2XSSC, 10% formamide and 0.1% Triton-X-100) for 5 times, 10 mins each. DAPI was dissolved in PBST and used for nuclei staining for 2 hours before the first cycle of sequencing. Finally, the sample was immersed in the washing and imaging buffer for imaging. Image acquisition was performed with Leica TCS SP8 confocal microscopy with 25X water-immersion objective (NA 0.95), with voxel size of 230 nm X 230 nm X 570 nm.

### 4. Data analysis

#### *In situ* sequencing analysis

A customized computational pipeline was built with *MATLAB (2019b)* to decode gene identity and quantify the gene expression level of each cell from the *in situ* sequencing images. First, sequencing fluorescence images were preprocessed with top-hat filtering by a disk structuring element (radius = 3) to remove the background noise. Second, the contrast of the image for each channel from the second to fifth sequencing cycle was adjusted to match the image from the first cycle with histogram matching function *“imhistmatchn”*. Third, the composite fluorescence images for the second to fifth cycle were registered with the composite fluorescence image from the first cycle using phase correlation algorithm followed by local distortion registration with function *“imregdemons”*. Fourth, the dots of amplicon locations were identified from images in the first cycle by a 3D regional maximum detection algorithm implemented in function *“imregionalmax”*. Then the dominant color of every identified dot in each cycle was determined by a 3×3×3 voxel volume surrounding its centroid location. The color sequence for each dot was decoded as a gene barcode and compared with the code-book. Fifth, cell segmentation was performed with ClusterMap^23^ or Stardist^25^ with custom cell mask dilation method, then RNA reads were assigned to the segmented cells accordingly.

*Python* package *Scanpy v1.6.0*^43^ was used for the single-cell gene expression analysis. Cells expressing less than 40 gene counts, expressing larger than 800 gene counts or only expressing less than three kinds of gene were filtered out. Gene counts of each cell were normalized so that the total count of all genes in each cell equals the median number of total counts across all cells. The normalized count value is then log-transformed with log(x+1). *Combat*^44^ was used to remove the potential batch effect among different imaging positions. Each gene in the cell-by-gene matrix was scaled to unit variance and zero mean followed by dimensionality reduction with principal components analysis (PCA). Based on the explained variance ratio, the top principal-components were used to construct the k nearest neighbor (kNN) graph for Leiden clustering^24^. Uniform Manifold Approximation and Projection (UMAP)^22^ was used to visualize the 2D representation of each cell. *Monocle 3* ^30^ is used to compute pseudotime along the cell trajectory.

#### Electrically recorded cell identification

Electrode position was located using the 3D electrode image collected by reflection-mode imaging and identified by the E-barcode positions. The electrode position in x and y coordinate was determined by the following steps: the electrode image was first projected to the x-y plane by maximum intensity projection (MIP) and transferred to gray-scale (pixel value ranging from 0-255). Then the MIP image was filtered with a global threshold of 50 to remove the non-electrode background. A 201-by-201 pixel size gaussian filter was applied to adaptively filter out the non-circular area, which is the I/O connect of the electrode. After locating the electrode in the x-y plane, the z coordinate of the electrode position was determined by fitting a 2D linear plane surface. The electrode recorded cell was further determined by calculating the area of intersection between each neighborhood CM cell and the electrode. The cell with the largest intersection area was identified as the electrically recorded cell.

#### Electrophysiology data processing

We followed the procedure of *SpikeInterface*^45^ to detect the spikes which passed the threshold (5*std of the signal) in one channel. Each spike has a fixed length of 1.6 second with the sampling rate of 10 kHz. After spike detection, the spikes were aligned at the minimum of the corresponding spike dv/dt, and then averaged to get a spike representation of that channel. For quality control, channels with less than two spikes within 60 seconds, spikes firing at the same time across multiple channels due to electrical shortage, or no electrical signals were excluded.

For each spike representation, we first denoised the spike with wavelet denoising using *PyWavelets*^46^. Then, the spike features were extracted through two cycles of down sampling operations. We first down sampled the whole 1.6 second length spike to 20 points (zoomed out binning) and then down sampled the 0.15 second length spike near the minimum of the differentiated spike to 42 points (zoomed in binning). In total, we generated 62 feature points for each spike representation of each channel.

#### Weighted Nearest Neighbor (WNN)

The weighted nearest neighbor (WNN)^29^, *i.e. FindMultiModalNeighbors* function from *Seurat v4,* was used to integrate the gene expression and electrophysiology data collected by *in situ* electro-seq. The principle component dimension for gene expression and electrophysiology was set as 7 and 6 (the elbow point in PCA variance), respectively. k=20 was used to find the k nearest neighbor and calculating the modality-specific weights. A WNN graph was then built as input to shared nearest neighbor joint clustering (Leiden)^24^, UMAP joint visualization and pseudotime derivation (*Monocle3*).

#### Gene expression, electrophysiology and WNN joint pseudotime analysis

We used a *R* package *Monocle3* for the pseudotime calculation of gene expression, electrophysiology features and the WNN joint representation described above. A set of hyperparameters (Euclidean distance ratio = 1, geodesic distance ratio = ⅔, minimal branch length = 5) in function *learn-graph* was used to first learn a principle graph of maturation. The node at the position of earliest stage was manually chosen as the root of principle graph to finalize the trajectory. Then the function *order_cells* was used to calculate the pseudotime.

#### Silhouette score

The Silhouette score was calculated by a *Python* package *silhouette_score* to quantitatively measure the separability of each cluster. The silhouette evaluates the quality of that clustering based on how well its data points are clustered. A silhouette measure is assigned to each data point representing how close a data point is to its own cluster in comparison to other clusters. The Silhouette score is calculated using the mean intra-cluster Euclidean distance (a) and the mean nearest-cluster Euclidean distance (b) for each sample. The Silhouette score for a sample is (b - a) / max (a, b). We calculated the Silhouette score for each stage pair (*e.g.*, Day12-Day46, Day12-Day21, etc.) in Extended Data Fig. 7 with the input of normalized data of electrically recorded CM gene expression, electrophysiology, WNN joint representation and all CM gene expression.

#### Sparse reduced-rank regression (RRR) model and bibiplot

For the RRR analysis^31^ we used 62 electrophysiological features across all 162 electrically recorded CMs. Both electrophysiological features and gene expression were normalized and z-scored as described above. Ninety-seven CM-correlated genes and ion channel genes were selected for the RRR analysis.

In brief, RRR finds a linear mapping of gene expression levels to a low-dimensional latent representation, from which the electrophysiological features are then predicted with another linear transformation. In Figure 4a-b, a model with rank r = 3, ridge penalty (α = 0.5), and lasso penalty (λ = 1.5) was used to yield a selection of 12 genes. In Extended Data Fig. 8e-f, a model with rank r = 5, ridge penalty (α = 0.5), and lasso penalty (λ = 1.5) was used to yield a selection of 32 genes. In Extended Data Fig. 8a-d, cross-validation was done by using 10 folds, elastic net α-values 0.5, 0.75, and 1.0, and λ-values from 0.2 to 6.0.

#### Coupled autoencoder prediction model

Coupled autoencoder^34–35^ was used for cross-modal prediction. Specifically, hyperparameter (latent loss weight =1, Adam optimizer with learning rate = 0.0001, batch size = 10, training epoch = 100, epoch step = 1000) and latent dimensionality (d =2) were used to capture the variability in the dataset. After training the coupled autoencoder network with the data collected in Day 12, Day 21, Day 46 and Day 64, continuously recorded electrophysiology data z-scored by the method mentioned above was used to infer continuous gene expression profiles for all the electrically recorded cells.

#### Data availability (for reviewers)

All the data are available in the Single Cell Portal (SCP, https://singlecell.broadinstitute.org/single_cell) with log in; password: electroseq123.

#### Code availability (for reviewers)

All the code are available in the GitHub repository (https://github.com/) with log in; password: electroseq123.

## Acknowledgements

We acknowledge the helpful discussions from J.S. and the early assistance from K.N., T.S. and Q.C. to this work. J.L. acknowledges the support from the Startup fund from the School of Engineering and Applied Sciences, Harvard University; NIH/NIMH 1RF1MH123948; NSF ECCS-2038603; Harvard Stem Cell Institute Collaborative Seed Grant SG-0124-20-00; Aramont Fund for Emerging Science Research; and the William F. Milton Fund. X.W. acknowledges the support from the Searle Scholars Program, Thomas D. and Virginia W. Cabot Professorship, and Edward Scolnick Professorship. Y.H. acknowledges the support from the James Mills Peirce Fellowship from the Graduate School of Arts and Sciences of Harvard University. Hao S. acknowledges the support from Aramont Fund for Emerging Science Research. Hailing S. acknowledges the support from a postdoctoral research fellowship from the Helen Hay Whitney Foundation.

## Author contributions

J.L. and X.W. conceived the idea. Q.L. cultured cells. R.L. fabricated and characterized mesh electronics. Q.L. and R.L. performed electrical recording. Q.L. and Z.L. prepared samples for *in situ* sequencing. Z.L. conducted *in situ* sequencing experiments. X.T., J.H., Y.H. and H.Z. analyzed data. All authors prepared figures and wrote the manuscript. J.L. and X.W. supervised the study.

## Additional information

**Correspondence and requests for materials** should be addressed to J.L. or X.W.

---

**Extended Data Fig. 1.**
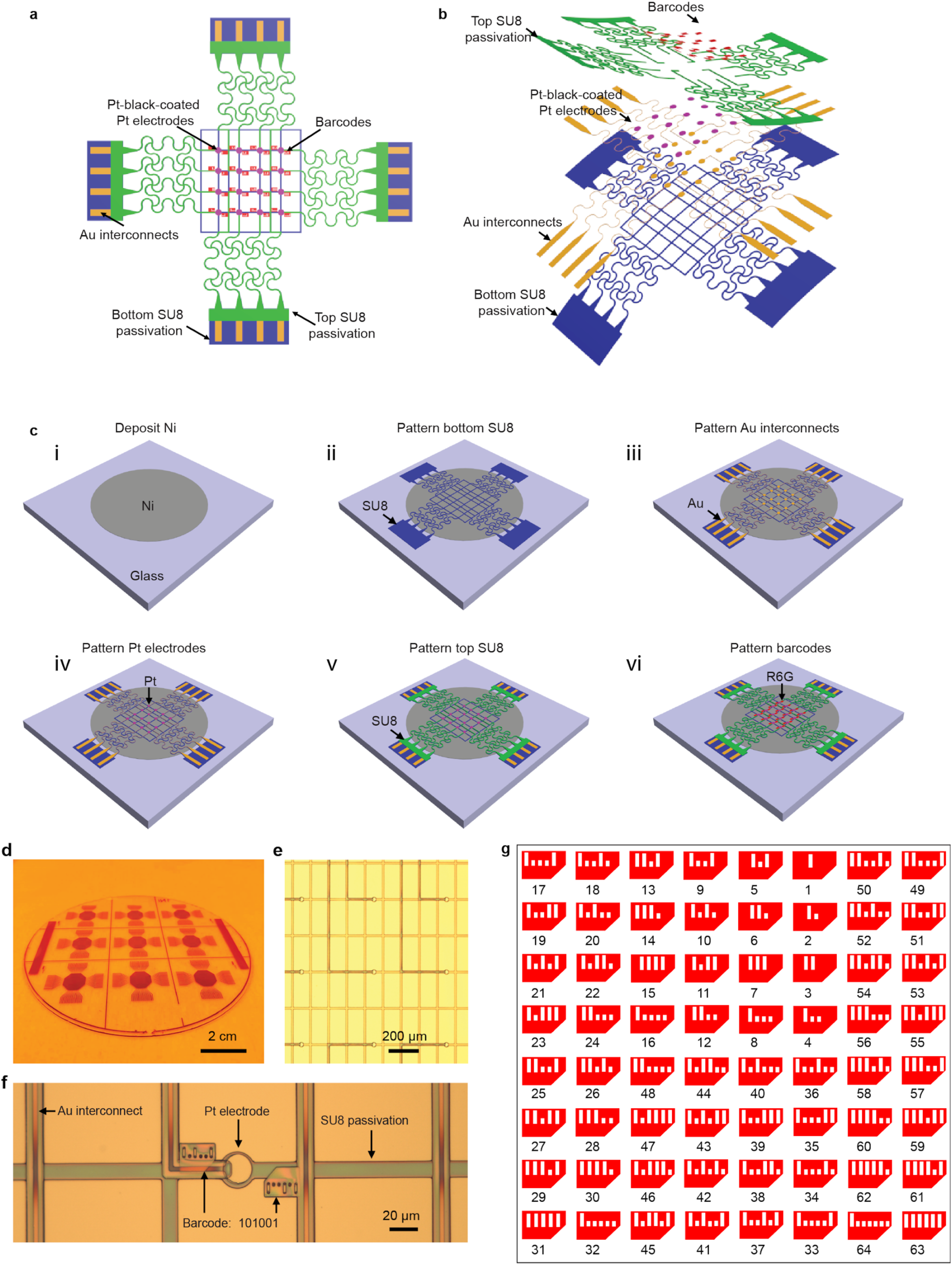
Design and fabrication of stretchable mesh electronics. **a**, Top-view schematic shows the structure of stretchable mesh electronics. **b**, Exploded-view schematic shows functional layers: Bottom SU-8 passivation layer (blue); Au interconnects (yellow); Pt black-coated electrodes (purple); top SU-8 passivation (green); and Electronic barcodes (E-barcodes) (red). **c**, Schematics show the key steps of fabrication flow of stretchable mesh electronics: (i) Deposition of 100-nm-thick Ni as the sacrificial layer on the glass substrate; (ii) Pattern of 400-nm-thick SU-8 as the bottom passivation layer; (iii) Pattern of 40-nm-thick Au as the interconnects; (iv) Pattern of 50-nm-thick Pt as the electrodes; (v) Pattern of 400-nm-thick SU-8 as the top passivation layer; (vi) Pattern of 400-nm-thick SU-8 doped with Rhodamine 6G as E-barcodes. **d**, Photograph of nine 64-channel stretchable mesh electronics fabricated on a 4-inch soda lime glass wafer. **e**. Bright-field (BF) optical image of an electrode array in stretchable mesh electronics on the substrate. **f**. BF optical image of a representative electrode with the paired E-barcodes and interconnects. **g**. Representative design of binary barcode for labeling 64 electrodes.

**Extended Data Fig. 2.**
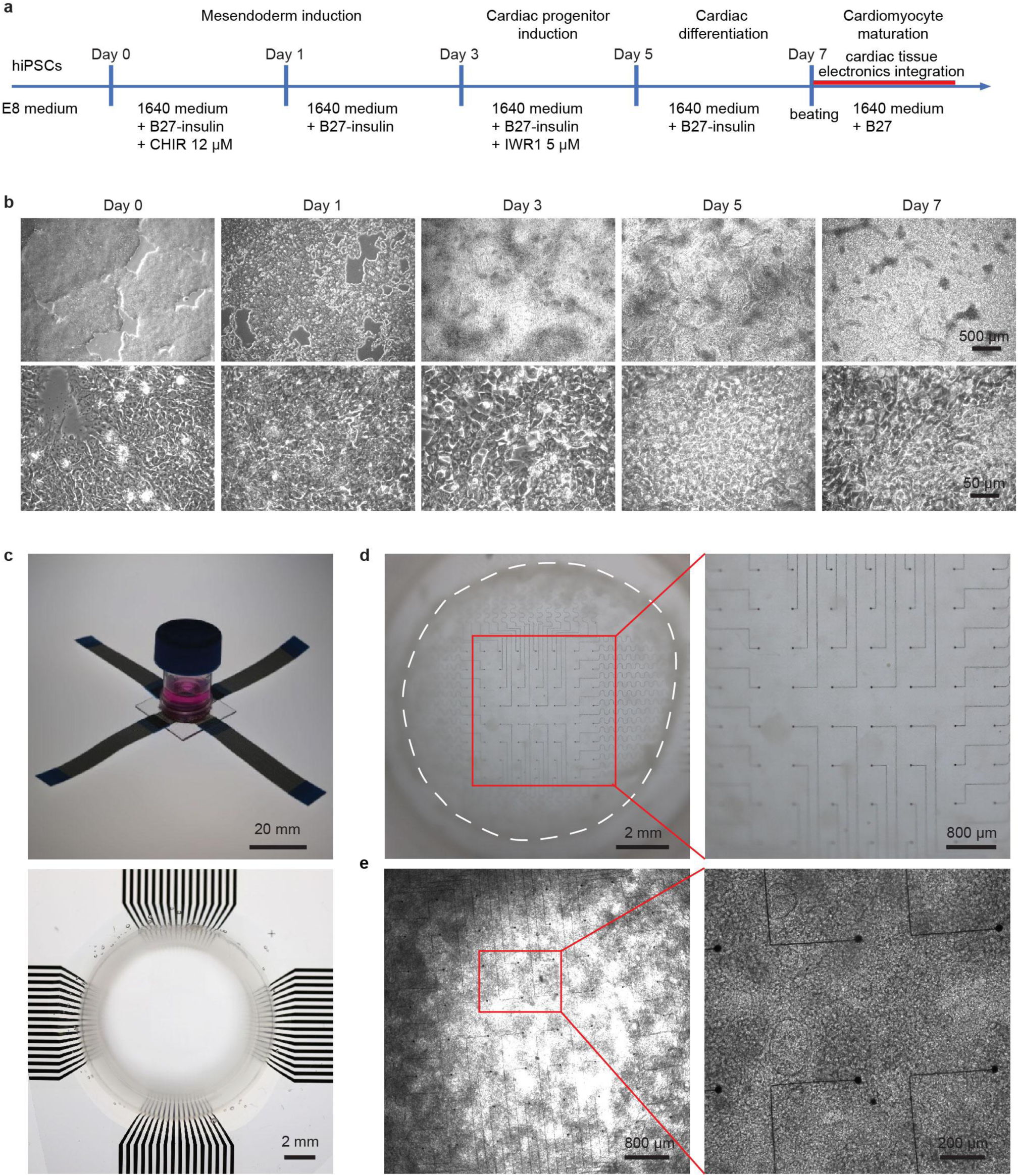
3D cardiac tissue-electronics hybrid. **a**, Schematics show the protocol for cardiac differentiation from human induced pluripotent stem cells (hiPSCs, IMR90-1) by canonical Wnt pathway signaling modulation with CHIR99021 and IWR1. The hiPSC-CMs will be dissociated into single cells and integrated with stretchable mesh electronics to form electronics-embedded cardiac patches. **b**, BF phase images show the cell morphology at Day 0, Day 1, Day 3, Day 5, and Day 7 of differentiation. **c**, Photographs show the side view (top) and top view (bottom) of the cell culture chamber bonded on the glass substrate with stretchable mesh electronics. **d**, Photographs show hiPSC-CM tissue-electronics hybrid in the cell culture chamber. The dashed line highlights the boundary of the hiPSC-CM tissue-electronics hybrid (left). Zoom-in BF image shows the cardiac tissue and stretchable mesh electronics from the red box (right). **e**, BF phase image shows the hiPSC-CM tissue-electronics hybrid (left). Zoom-in image highlights the interaction between hiPSC-CM tissue and stretchable mesh electronics (right).

**Extended Data Fig. 3.**
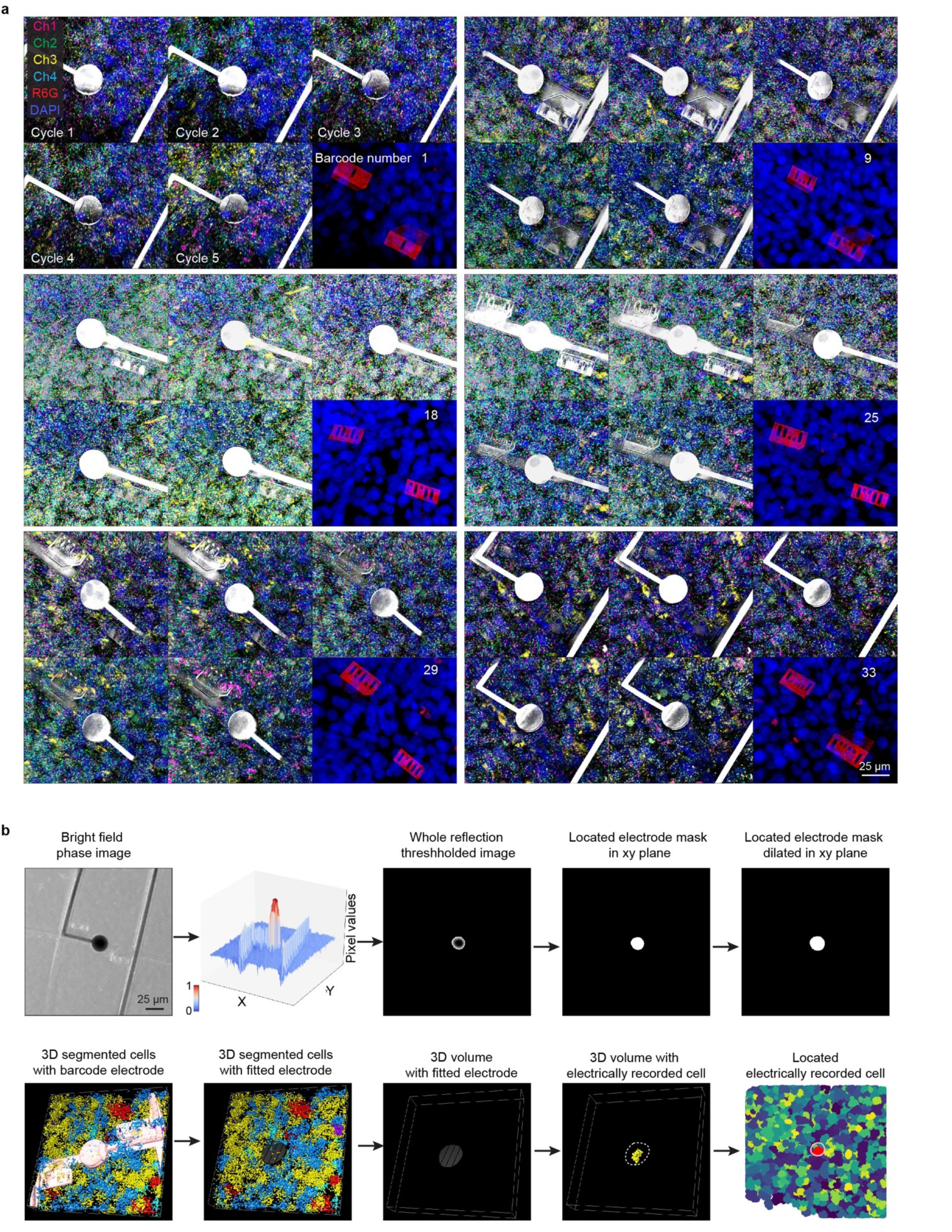
Locating of electrically recorded CMs during 3D *in situ* electro-seq. **a**, Six representative raw fluorescence images of five cycles sequencing and one cycle E-barcode imaging at Day 46. Ch, color code for the four fluorescence channels; Electrodes were imaged using reflection mode and highlighted as grey; E-barcodes were labelled with the fluorescent dye Rhodamine G (R6G); DAPI, cell nuclei staining. **b,** Schematics of finding electrically recorded CM. Bright field image was projected to the x-y plane and transferred to a gray-scale image, which was gaussian filtered and dilated to find the electrode mask in the x-y plane. The electrode mask in 3D space was fitted as a linear 2D surface. The electrically recorded cell was identified as the cell having the most intersection with the electrode mask in 3D space.

**Extended Data Fig. 4.**
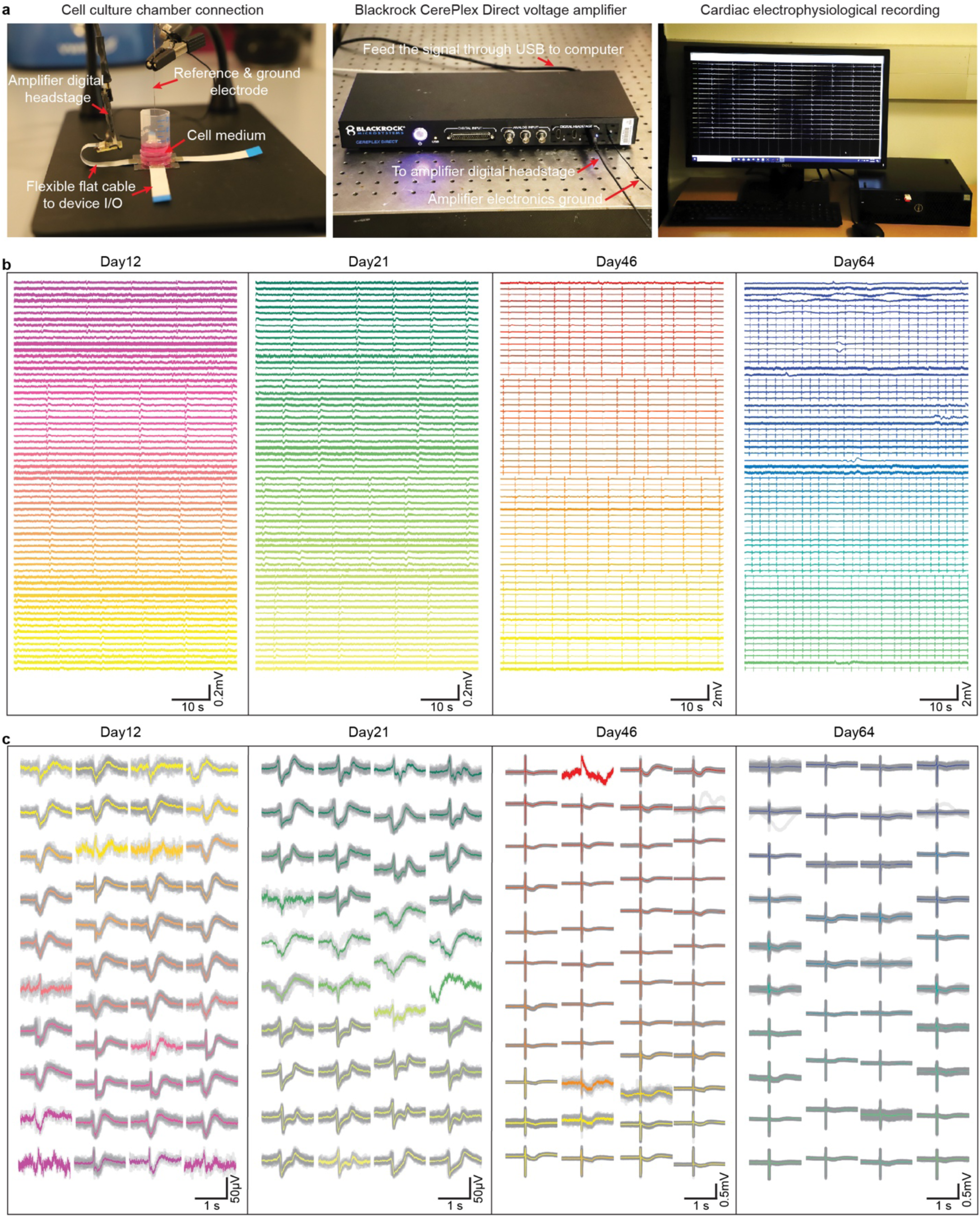
Electrophysiological mapping of hiPSC-CM tissues over the time course of *in vitro* maturation. **a**, Photographs show the multiplexing recording setup that measures hiPSC-CM tissues electrophysiology. **b**, Representative 64-channel voltage traces recorded from the hiPSC-CM tissues at Day 12, Day 21, Day 46, and Day 64 of differentiation, respectively. **c**, Representative single-spike action potentials recordings at Day 12, Day 21, Day 46, and Day 64 of differentiation, respectively.

**Extended Data Fig. 5.**
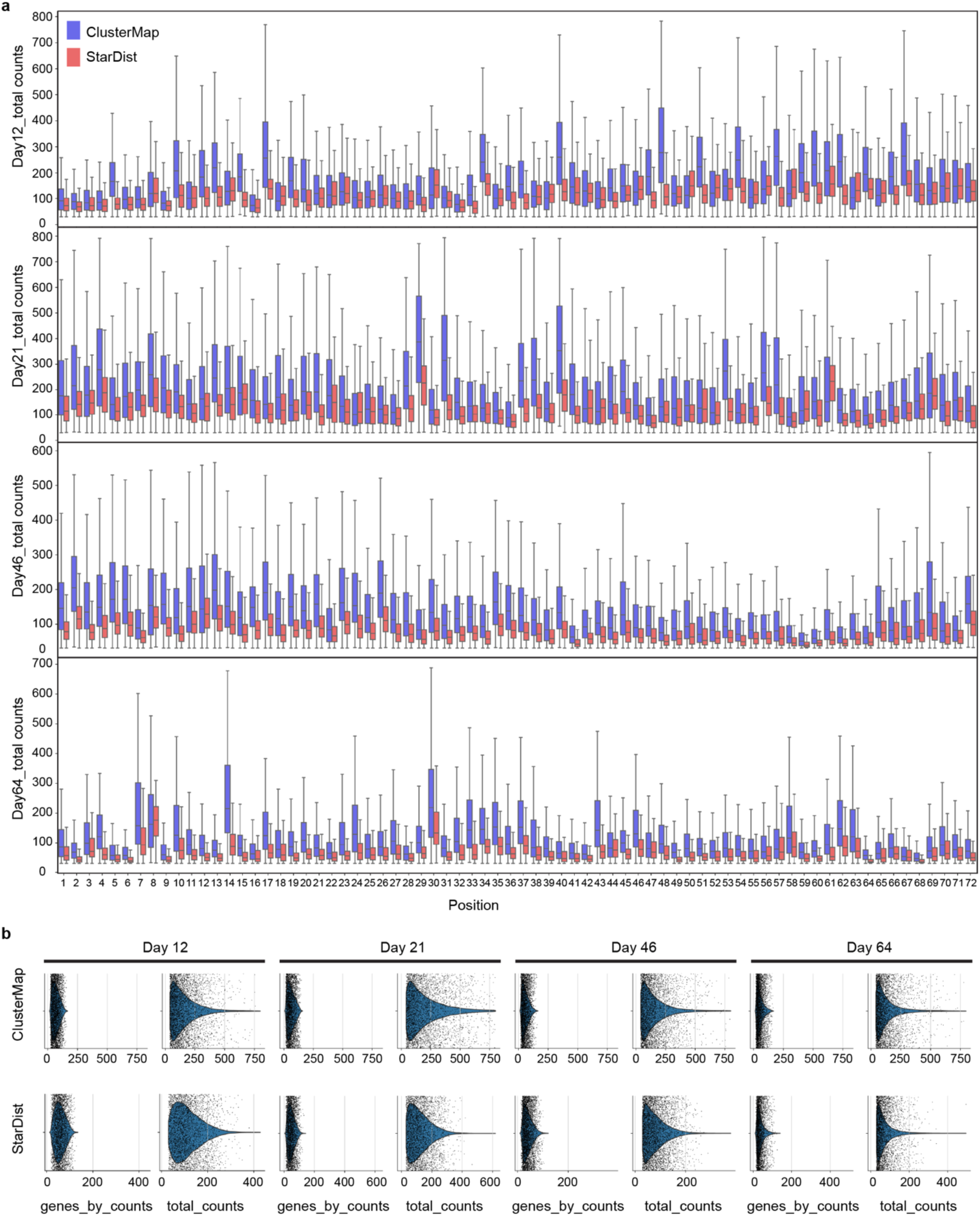
Comparison of cell segmentation performance by ClusterMap and StarDist methods. **a**, Box plots show the total RNA counts per cell in each position of cardiac tissues at Day 12, Day 21, Day 46, and Day 64 of differentiation, respectively, using ClusterMap and StarDist cell segmentation methods. 72 positions were imaged and analyzed for each sample. **b**, Violin plots show the distribution of total RNA counts and gene counts per cell at Day 12, Day 21, Day 46, and Day 64, of differentiation respectively, using ClusterMap and StarDist cell segmentation methods. The results show that ClusterMap can identify more RNA counts per cell from our samples compared with the SarDist method.

**Extended Data Fig. 6.**
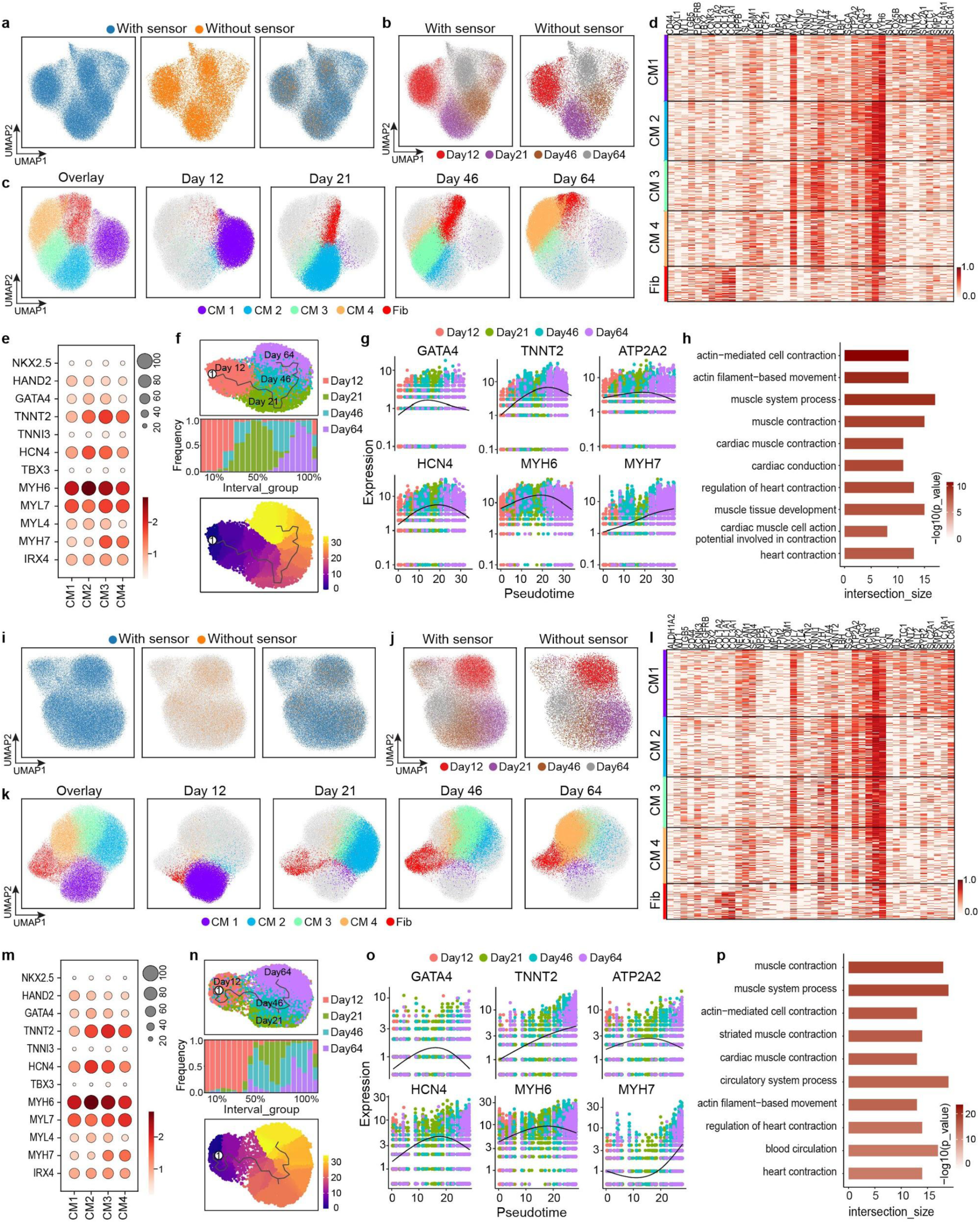
Comparison of cell typing analysis by ClusterMap and StarDist methods. Cell typing information in **a-h** and **i-p** were from ClusterMap- and StarDist-based cell segmentations, respectively. **a** and **i**, UMAP visualizations of all the cells from electronics-embedded cardiac tissues and control cardiac tissues (without electronics embedding) show the similar distribution, which proves negligible effects from electronics embedding on the gene expression. **b** and **j**, UMAP visualizations of all the cells from cardiac electronics-embedded cardiac tissues and control cardiac tissues (without electronics embedding) labeled by the different days of differentiation show the similar cluster distributions, which proves negligible effects from electronics embedding on the gene expression over maturation. Colors correspond to days of differentiation. **c** and **k**, UMAP visualizations highlight the cell types clustered by Leiden clustering and their distributions at Day 12, Day 21, Day 46, and Day 64 of differentiation, respectively. Colors correspond to different cell types. **d** and **l**, Heatmaps of top 42 differentially expressed genes aligned with each cell type. Colors correspond to the normalized gene expression value. The values are normalized to 0 to 1 for each gene. **e** and **m**, Dot plots of selected marker genes expressions in cardiac tissues. The size of the dot corresponds to the percentage of cells within a cell type, and its color corresponds to the average expression level. **f** and **n**, Top: UMAP visualizations show the trajectory of the cardiac tissue maturation using only CM cells. Colors correspond to days of differentiation. The line corresponds to the principal graph learned by Monocle 3. Middle: Stacked bar plot shows percentage of cells across inferred pseudotime of the cardiac tissue development. Colors correspond to days of differentiation. Bottom: UMAP visualizations show the trajectory of cardiac tissue maturation. Colors correspond to inferred pseudotime. The line corresponds to the principal graph learned by *Monocle 3*. The trajectory starting anchor was manually chosen on the graph position of Day 12 as the start of the pseudotime. **g** and **o**, Kinetics plots shows the relative expression of cardiac lineage marker genes across maturation pseudotime. Colors correspond to days of differentiation. **h** and **p**, Gene Ontology analysis shows significant terms related to cardiac muscle contraction, conduction, and development. Bar plot displays the top 10 significant (FDR < 0.05) gene ontology (GO) terms enriched in electrophysiological related genes, mostly involved in cardiac muscle contraction, conduction, and development.

**Extended Data Fig. 7.**
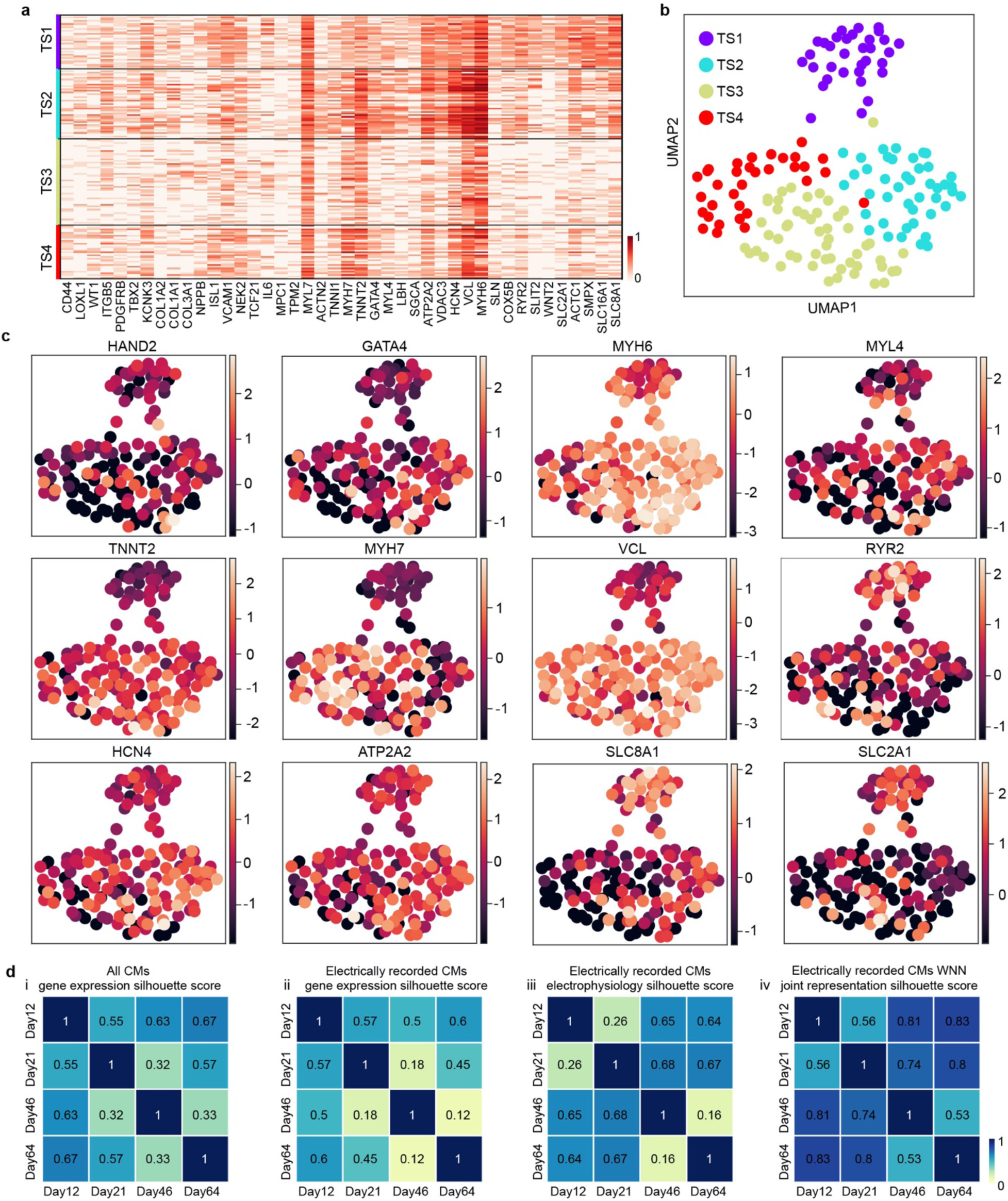
Clustering results of electrically recorded CMs by *in situ* electro-seq. **a**, Heatmap for top 42 differentially expressed genes aligned with each transcriptional state (TS). The values are normalized to 0 to 1 for each gene. **b**, UMAP visualizations highlight the cell types clustered by Leiden clustering. Colors correspond to different TSs. **c**, UMAP visualization shows CM cell type-related genes expression across all the electrically recorded CMs. Colors correspond to z-scored expression level. **d**, Matrix of silhouette scores measuring the separability among Day 12, Day 21, Day 46, and Day 64 of differentiation: (i) All CMs gene expression; (ii) electrically recorded CMs gene expression; (iii) electrically recorded CMs electrophysiology; and (iv) electrically recorded CMs WNN joint representations from gene expression and electrophysiology.

**Extended Data Fig. 8.**
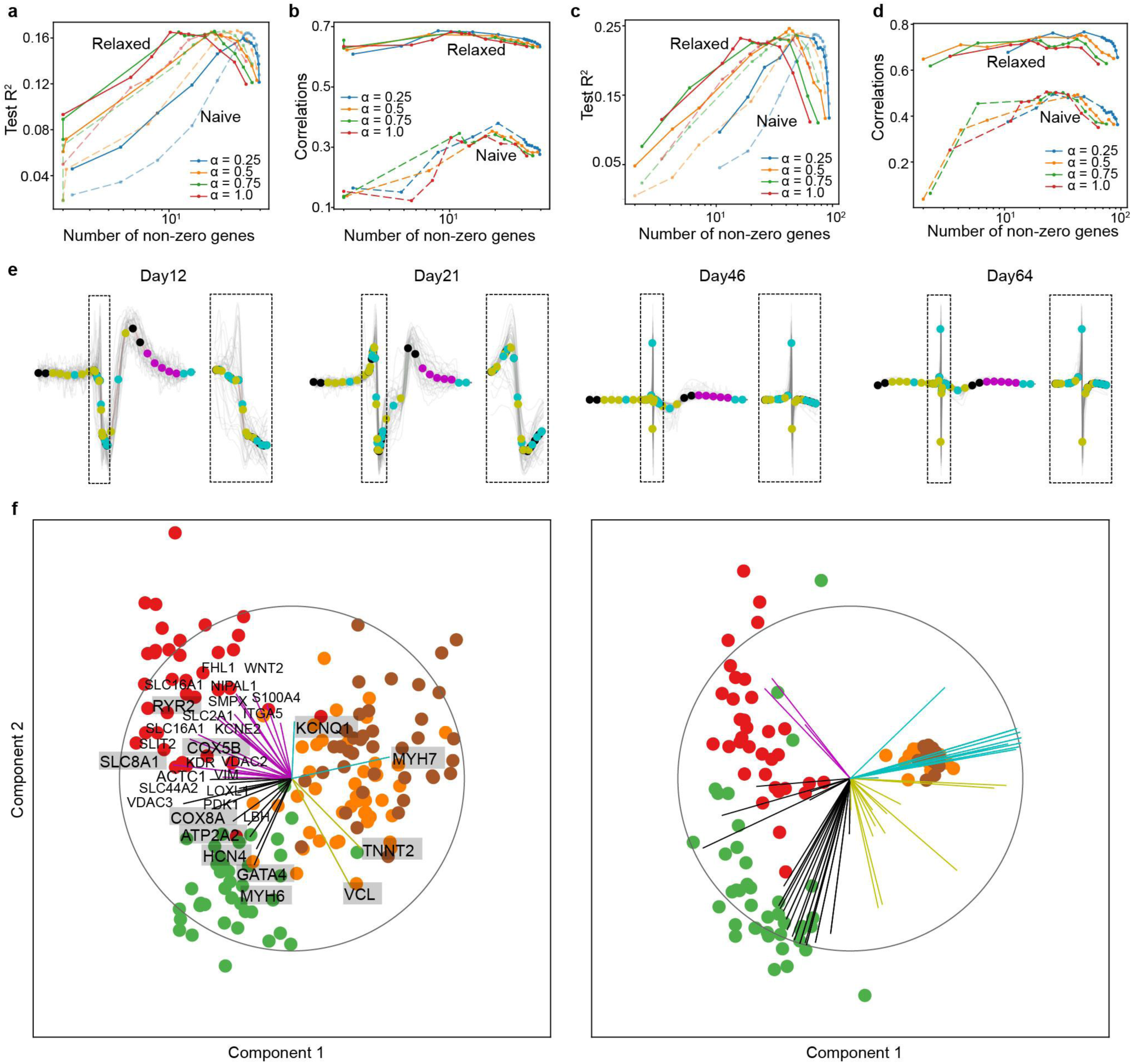
Cross-modal visualization and correlation of *in situ* electro-seq results of hiPSC-CM maturation by reduced-rank regression (RRR) analysis. **a,** Test R^2^ of ‘relaxed’ and ‘naive’ sparse RRR for ion channel related genes with α = 0.25, 0.5, 0.75, and 1, respectively. **b**, Cross-validated correlations of ‘relaxed’ and ‘naive’ sparse RRR for ion channel related genes with **α** = 0.25, 0.5, 0.75, and 1, respectively. **c,** Test R^2^ of ‘relaxed’ and ‘naive’ sparse RRR for 97 CM-related top differentially expressed genes with α = 0.25, 0.5, 0.75, and 1, respectively. **d**, Cross-validated correlations of ‘relaxed’ and ‘naive’ sparse RRR for 97 CM-related differentially expressed genes with α = 0.25, 0.5, 0.75, and 1, respectively. **e,** Representative 62 electrophysiological features are extracted through down sampling operations for each spike waveform on Day 12, Day 21, Day 46, and Day 64 of differentiation, respectively. 1.6-second waveforms are sampled to 20 bins. Inset: 0.15-second fast spikes are sampled to 42 bins. **f**, A sparse RRR model to visualize and align t-states and e-states. Components 1 and 2 of the rank-5 model are shown, n=162. The model selects 97 and 32 most related genes to train the model and visualize, respectively.

**Extended Data Fig. 9.**
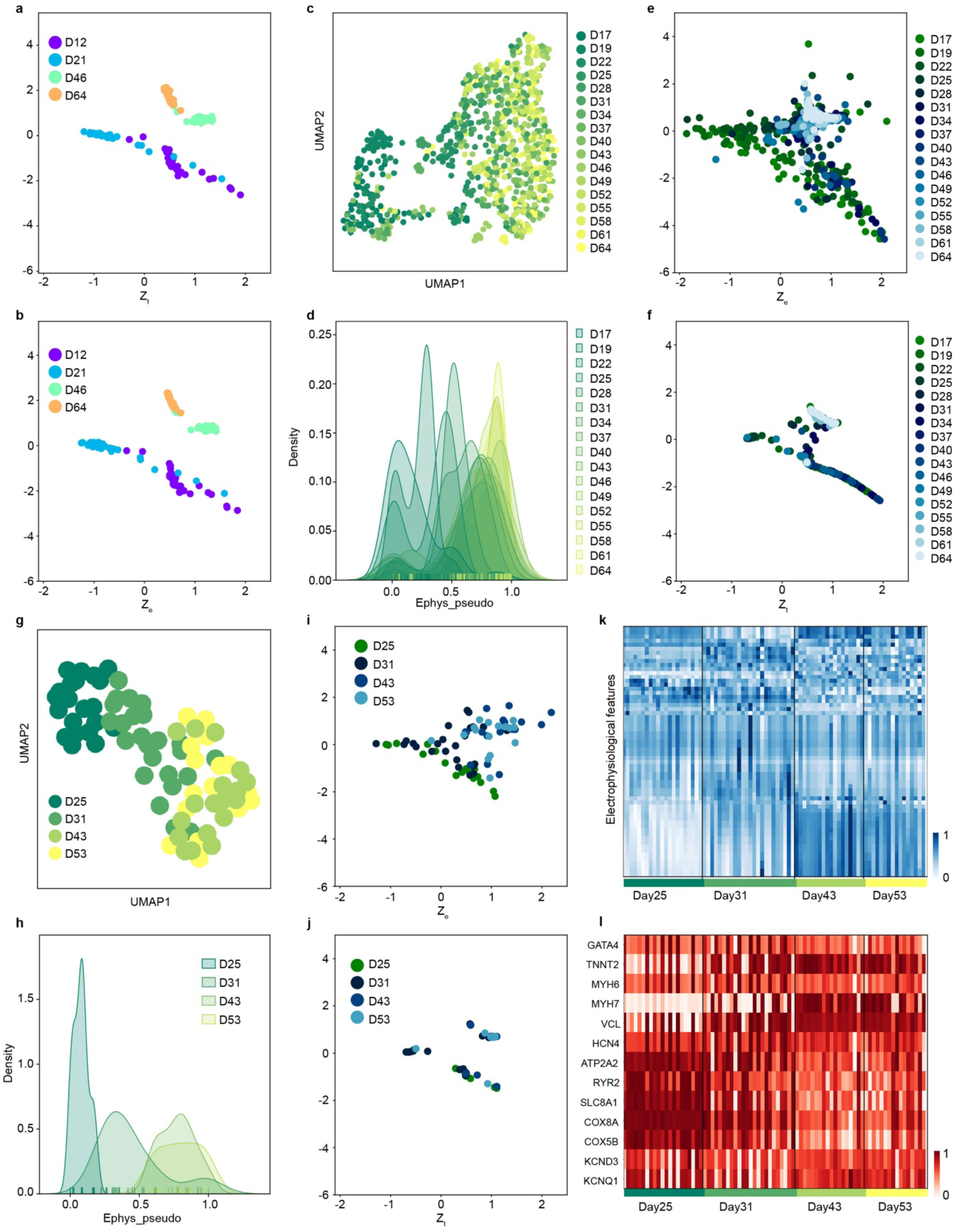
Cross-modal prediction of *in situ* electro-seq results of hiPSC-CM maturation by coupled autoencoder. **a-b**, UMAP visualizations of the extracted features of single spike waveforms from the continuous electrical recording of one hiPSC-CM patch is color coded by days of differentiation (**a**) and pseudotime by *Monocle3* normalized to 0-1 (**b**). **c-d**, Distribution plots shows pseudotime distributions by *Monocle3* using electrophysiology color coded by days of differentiation. **e**, Coupled autoencoder-encoded 2D representations trained by *in situ* electro-seq data show the distribution of transcriptional (*Zt*) and electrophysiological (*Ze*) data. Colors correspond to days of differentiation. **f**, Coupled autoencoder-encoded 2D representation shows cross-modal predicted gene expressions (*Ze-t*) from continuous electrical recording using the trained coupled autoencoder. **g-h**, Distribution plots show pseudotime distributions by *Monocle3* using electrophysiology color coded by days of differentiation for the 3D cardiac organoid. **i**, Coupled autoencoder-encoded 2D representation shows continuous electrical recording (*Ze*) using the trained coupled autoencoder for the 3D cardiac organoid. **j**, Coupled autoencoder-encoded 2D representation shows cross-modal predicted gene expressions (*Ze-t*) for the 3D cardiac organoid. **k**, Heatmap shows single-cell electrophysiological features predicted for the 3D cardiac organoid. The values are normalized to 0 to 1 for each feature. **l**, Heatmap shows single cell resolution gene expression predicted for the 3D cardiac organoid. 13 genes were selected according to the RRR plot results. The values are normalized to 0 to 1 for each gene.

**Extended Data Fig. 10.**
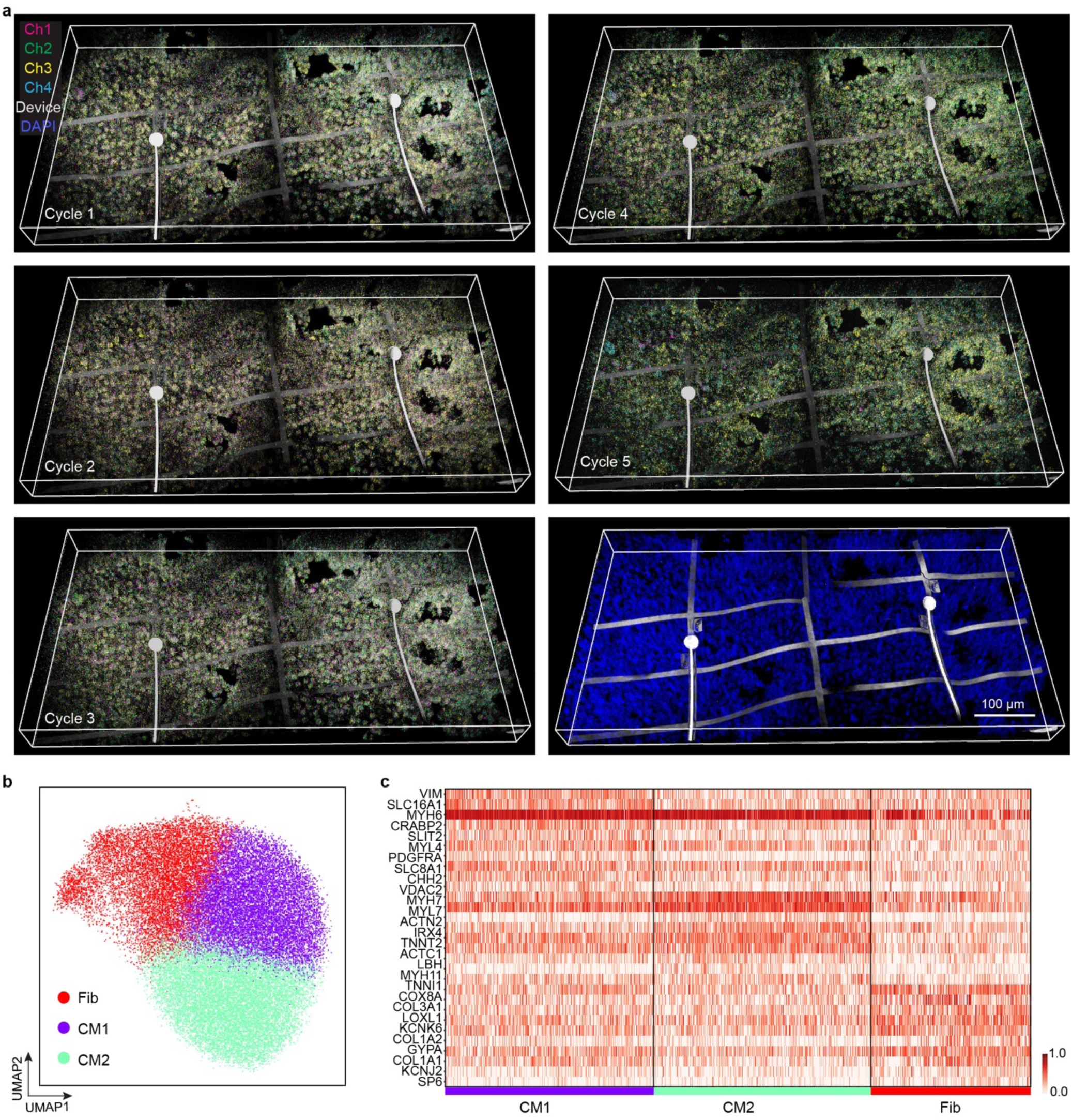
*In situ* electro-seq of cardiac tissue with heterogeneous CM types. **a**, Raw fluorescence images of five cycles of *in situ* sequencing and one cycle of barcode imaging. Ch, color code for the four fluorescence channels; Electrodes were imaged using reflection mode and color coded as grey; DAPI, cell nuclei staining. **b**, UMAP visualizations of all the cells from the heterogeneous cardiac tissue show three types of cells, fibroblast (Fib), cardiomyocytes 1 (CM1), and CM 2. **c**, Heatmap of 28 top differentially expressed genes aligned with each cell type. Color corresponds to normalized expression value.

